# Biased Signaling: Distinct Ligand-directed Plasma Membrane Signalosomes Using a Common RGS/G protein Core

**DOI:** 10.1101/666578

**Authors:** Timothy J. Ross-Elliott, Justin Watkins, Xiaoyi Shan, Fei Lou, Bernd Dreyer, Meral Tunc-Ozdemir, Haiyan Jia, Jing Yang, Luguang Wu, Yuri Trusov, Patrick Krysan, Alan M. Jones

**Author notes:** Correspondence: Dr. Alan M. Jones, Address: Department of Biology, The University of North Carolina at Chapel Hill, Coker Hall, CB#3280. Abbreviations: *Arabidopsis thaliana* Regulator of G Signaling (AtRGS1), Clathrin-Mediated Endocytosis (CME), Sterol-Dependent Endocytosis (SDE), Total Internal Reflection Fluorescence Microscopy (TIRF), Clathrin Light Chain (CLC), Flotilin 1 (FLOT1), Vacuolar Protein Sorting 26 (VPS26).

## Abstract

Biased signaling occurs when different ligands that are directed at the same receptor launch different cellular outcomes. Because of their pharmacological importance, we know the most about biased ligands and little is known about other mechanisms to achieve signaling bias. In the canonical animal G protein system, endocytosis of a 7-transmembrane GPCR is mediated by arrestins to propagate or arrest cytoplasmic signaling depending on the bias. In Arabidopsis, GPCRs are not required for G protein coupled signaling because the heterotrimeric G protein complex spontaneously exchanges nucleotide. Instead, the prototype 7-transmembrane Regulator of G Signaling 1 protein AtRGS1 modulates G signaling and through ligand-dependent endocytosis, de-repression of signaling is initiated but canonical arrestins are not involved. Endocytosis initiates from two separate pools of plasma membrane: sterol-dependent domains, possibly lipid rafts, and a clathrin-accessible neighborhood, each with a select set of discriminators, activators, and newly-discovered arrestin-like adaptors. Different trafficking origins and trajectories lead to different cellular outcomes. Thus, compartmentation with its attendant signalosome architecture is a previously unknown mechanism to drive biased signaling.

---

Different ligands discriminated by the same receptor or utilizing the same core of signaling elements can set in motion a cascade of events that lead to multiple outcomes. When subsets of outcomes are ligand-specific, we label this biased signaling. This bias can be encoded in the ligand structure (biased ligands), in the receptors (biased receptors), or in the interactions between the signaling components (biased systems) ^1,2^. Of the three defined mechanisms, biased ligands is the most widely studied because of its immediate pharmacological significance, such as the development of drugs that provide analgesia without the addictive side effect ^3–5^. **a)** <u>Ligand bias</u> occurs when a ligand stabilizes one or a subset of conformations of a receptor protein which then preferentially recruits or activates signaling elements such as arrestin and the heterotrimeric G protein complex that lead toward one cellular outcome over another ^6,7^. For example, some ligands are biased towards β-arrestins (aka arrestin-2 and −3) signaling ^8–10^ such as certain dopamine agonists compared to dopamine which is biased towards D_2_R-β-arrestin coupling ^11^. Certain opioid agonists compared to morphine are biased toward G-protein-coupled signaling ^4,7,12^. **b)** <u>Receptor bias</u> occurs through recognition of the same ligand by multiple receptor types ^13^ including so-called decoy receptors ^14,15^, some of which may in fact be functional receptors that use non-classical signaling pathways ^14,16^. Dopamine D_1_ and D_2_ class receptors recognize dopamine, but signal through different subunits of Gα as well as arrestin ^17^. **c)** <u>System bias</u> involves a cell-mediated shift to one pathway over another by some unknown mechanism but one possibility is through mass action, for example, making arrestin more abundant at the receptor than Gα or a particular kinase ^18^. This last category of bias signaling is the least understood and the subject of the present work.

Phosphorylation patterns are the chemical bar codes for biased signaling at least for arrestin-dependent outcomes ^19–23^. G-protein coupled Receptor Kinases (GRKs) phosphorylate the C-terminus or other cytoplasmic elements of GPCRs in response to agonist binding ^24^. The subsequent coupling of arrestin to phosphorylated GPCRs does not end G-protein-independent signaling. Arrestin-bound GPCRs further activates some kinase pathways including MAPK and tyrosine kinases ^25–27^ but not all (e.g. ref. ^28^). While plants lack GRKs, an Arabidopsis family of kinases called WITH NO LYSINE (**WNK**) kinase phosphorylate the C-terminal tail of the non-canonical 7-transmembrane Regulator of G Signaling (**AtRGS1**) in response to extracellular glucose ^2,29^. AtRGS1 is also phosphorylated by other kinases including BAK1, a co-receptor for flg22 which is a Pathogen-Associated Molecular Pattern (**PAMP**) ^30–32^ and loss of AtRGS1 and other G protein components severely affect resistance in a wide range of pathogens ^33^. System-biased signaling may be particularly relevant in plants where AtRGS1 modulates intracellular signaling transduction through G protein activation. AtRGS1 regulates G protein activation with ligand discrimination likely facilitated by membrane bound Receptor Like Kinases (RLKs) of which there are more than 400 members in Arabidopsis ^34^.

In animals, activation of G Protein signaling results from GDP exchange for GTP on the Gα subunit, wherein this nucleotide exchange is the rate-limiting step ^35^. Desensitization of the cell toward the signal occurs through endocytosis of the GPCR mediated by arrestins ^36,37^. In Arabidopsis, however, the Gα subunit, AtGPA1, spontaneously exchanges nucleotides without a GPCR thus it is self-activating with the intrinsic GTPase activity being the rate-limiting step ^38,39^. AtRGS1 accelerates the intrinsic GTPase of AtGPA1 ^40^ and as a result, inactive Gα remains bound to GDP until de-repression through AtRGS1 endocytosis, permitting Gα activation through nucleotide exchange and subsequent downstream signal transduction ^29^. This AtRGS1 endocytosis is a well-established proxy for sustained G protein activation and the proportion of endocytosed AtRGS1 is linearly related to the proportion of Gα in the GTP state ^2^. In Arabidopsis, sugars ^2,41^ and flg22 ^32^ activate AtRGS1 ^29^.

Physically de-coupling AtRGS1 from the heterotrimeric G protein complex by endocytosis is the engine of the de-repression mechanism, at least for sustained activation. There are two modes of endocytosis in plants: Clathrin-Mediated Endocytosis (**CME**) and Sterol-Dependent Endocytosis (**SDE**) ^42^, however, the protein that directly couples to AtRGS1 and is responsible for endocytosis is unknown. Extensive work elucidated the CME pathway in animal and yeast systems with the function of many molecular components being well characterized. Of particular interest is the ADAPTOR PROTEIN COMPLEX-2 (**AP-2**) that is required for recognition and binding of cargo ^43–45^, recruiting clathrin to the PM, and the subsequent formation of clathrin-coated vesicles. In the absence of AP-2 function, clathrin-coated vesicle formation is reduced and endocytosis is inhibited ^46^. The AP-2 complex is a heteromeric complex consisting of 2 large subunits (α2 and β2), 1 medium subunit (µ2), and 1 small subunit (σ2) ^47^.

SDE is a clathrin-independent mechanism for internalization of membrane-associated proteins. Sometimes referred to as lipid raft endocytosis, SDE of these microdomains requires flotilin1 (**Flot1**), and possibly the microdomain-associated remorin protein ^48^ for internalization of sterol-rich vesicles ^49^. Membrane proteins PIP2;1 and Respiratory Burst Oxidase Homolog D (**RbohD**) are selectively internalized via sterol-dependent endocytosis under salt stress conditions ^50,51^.

Here, we present data illustrating a biased system where two distinct extracellular ligands induce endocytosis of AtRGS1 from separate plasma membrane origins, of which one is mediated by a an arrestin-fold-containing protein, Vacuolar Protein Sorting 26 (VPS26). flg22 initiates AtRGS1 endocytosis via CME, while glucose activates both pathways of endocytosis. Phosphorylation of AtRGS1 and involvement of individual subunits of the heterotrimeric G protein complex are also ligand specific as are the immediate downstream consequences. From the CME-mediated-AtRGS1 origin, flg22 induces a MAPK cascade known to drive transcriptional reprogramming ^52^, whereas from the SDE-mediated-AtRGS1 origin, glucose induces transcriptional change that is independent of the MAPK cascade.

## Results

### G signaling activation via AtRGS1 endocytosis by two distinct extracellular signals

To date, there are 2 well-studied signals that directly and rapidly activate the Arabidopsis heterotrimeric G protein signaling pathway through AtRGS1 endocytosis: **1)** flg22 which is a 22-amino acid PAMP from the plant pathogen *Pseudomonas syringe* that is recognized by plant cells to initiate the innate immunity pathway ^53–55^. It is already established that flg22 is perceived extracellularly (e.g. ref. ^56^ by co-receptors BAK1 and FLS2 as part of a larger G protein complex ^57,58^). **2)**. Glucose or a glucose metabolite which are products of photosynthesis ^59^ and metabolism of starch stores ^60^. An example of a time course for activation by flg22 and D-glucose is shown in Figure 1A.

**Figure 1.**
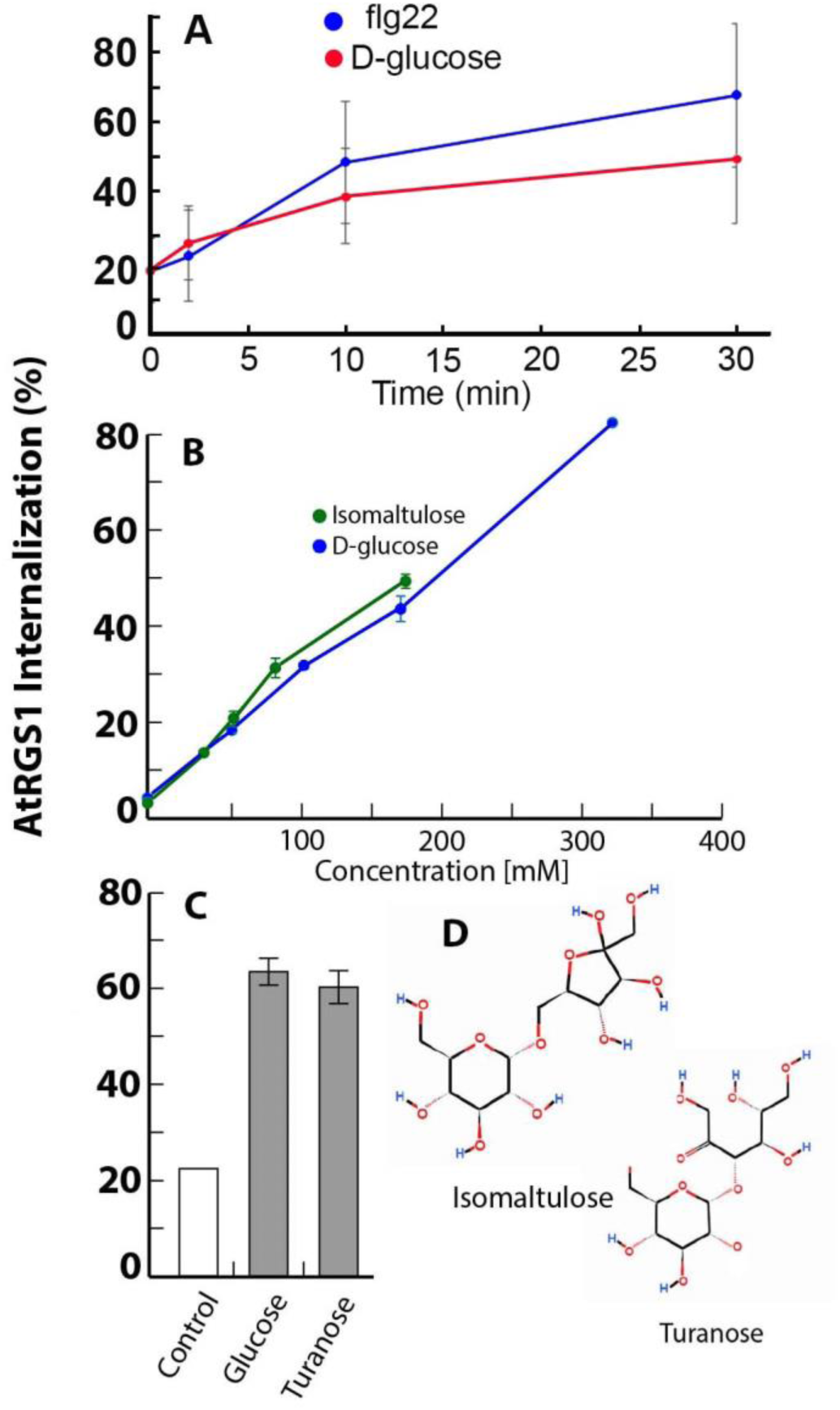
An AtRGS1 complex perceives extracellular flg22 and glucose or glucose metabolite. **A**. flg22 and D-glucose induce rapid endocytosis of AtRGS1-YFP. Values represents the amount of AtRGS1 that is internal to the cell as a percent of the total YFP fluorescence. The type of raw data that is used to generate these values is shown in Supplemental Figure S1C. **B**. While flg22 is perceived extracellularly, an extracellular site of glucose perception for activation of G signaling in Arabidopsis has not yet been made with an impermeant glucose analog. Despite being 9-fold less permeant to the Arabidopsis plasma membrane, isomaltulose is as or slightly more potent than D-glucose in activation of G signaling as measured by proxy using AtRGS1-YFP endocytosis ^2^. See supporting data for this panel in Supplemental Material Figure S1. **C**. Turanose is impermeant to plant cells ^69^ yet is as effective as D-glucose in inducing AtRGS1-YFP endocytosis. The purity of turanose was >98%. **D**. Structures. Both isomaltulose and turanose share a glucose ring moiety.

Many sugars, although primarily sucrose, are transported extracellularly in the apoplastic space where they are converted to glucose by cell wall localized invertases and potentially further metabolized to a signal. These sugars are rapidly taken up through a large family of transporters ^61,62^ where they are detected in the cytoplasm, but some sugars may also be detected extracellularly ^63^. D-glucose and some other glucose containing sugars induce rapid endocytosis of AtRGS1-dependent sugar signaling ^2,29^ and because AtRGS1 shares the membrane topology of GPCRs which perceive extracellular signals, it has been assumed that the extracellular glucose or metabolite is perceived by the AtRGS1/G protein complex. However, neither an extracellular site for perception nor direct evidence for agonist binding to AtRGS1 has been shown. To address the former, impermeant sugars were tested for the ability to activate G signaling.

The glucose-fructose dissacharide 6-0-α-D-glucopyranosyl-D-fructose (aka isomaltulose, pallatinose), is presumed not to be transported across the plant cell membrane although it acts as an active glucose precursor if synthesized intracellularly ^64,65^. While the expression of sucrose isomerase in potato increased apoplastic isomaltulose, transport across any membrane has yet to be demonstrated ^66^. Importantly, isomaltulose is not hydrolyzed extracellularly ^67,68^. To determine if isomaltulose is impermeant to the plasma membrane, we chemically synthesized [^14^C] isomaltulose (Figure S1A) and tested for uptake into Arabidopsis seedlings. Isomaltulose was at least 9-fold less permeant to cells than glucose (Figure S1B). Therefore, to determine if sugars are perceived extracellularly by hypocotyl cells, we tested the effect of isomaltulose on AtRGS1-YFP endocytosis (Figure S1C). Whereas several mono and disaccharides failed to activate G signaling, extracellular isomaltulose caused AtRGS1 internalization more effectively (P< 0.01) than D-glucose (Figure 1B) despite being transported ∼10-fold less suggesting that isomaltulose activates AtRGS1 extracellularly. The 5% D-glucose impurity in the isomaltulose preparation would induce at most 5% AtRGS1 internalization and therefore does not account for the observed level of activation. Turanose is another disaccharide that is thought to be perceived extracellularly ^64^ and is impermeant ^69^ therefore endpoint analysis was performed using this sugar, and just as for both glucose and isomaltulose, turanose activated G signaling (Figure 1C). Isomaltulose and turanose are disaccharides that share a glucose ring (Figure 1D), suggesting that glucose or a glucose metabolite is the discriminated signal or is important for a metabolic agonist (e.g. sugar nucleotides).

### Biased signaling outputs

Two rapid events of the flg22 response is the induction of MITOGEN ACTIVATED PROTEIN KINASE 6 (MPK6) activity and Ca^2+^ signaling ^70^. To test the impact of flg22 and D-glucose on MPK6 activity in etiolated hypocotyls, we developed a FRET-based sensor that measures kinase activity specifically for MPK6, called *S*ensor *O*f *M*APK *A*ctivity (SOMA) ^71^. The SOMA lines were tagged with either the human immunodeficiency virus 1 (HIV-1) nuclear export signal (SOMA-NES) or the SV40 nuclear localization signal (SOMA-NLS) ^72,73^ to measure MPK6 activity in the cytosol or nucleus, respectively. Validation of these reporters for hypocotyl epidermal cells using positive and negative controls is shown in Supplemental Figure S2A-F. As shown in Figures 2A-F, rapid FRET gains were observed in both SOMA-NES and SOMA-NLS within 2-4 minutes after treatment with flg22. When treated with 6% D-glucose, no change in FRET efficiency was observed, suggesting that D-glucose does not induce activity of MPK6 (Fig 2G and H).

**Figure 2.**
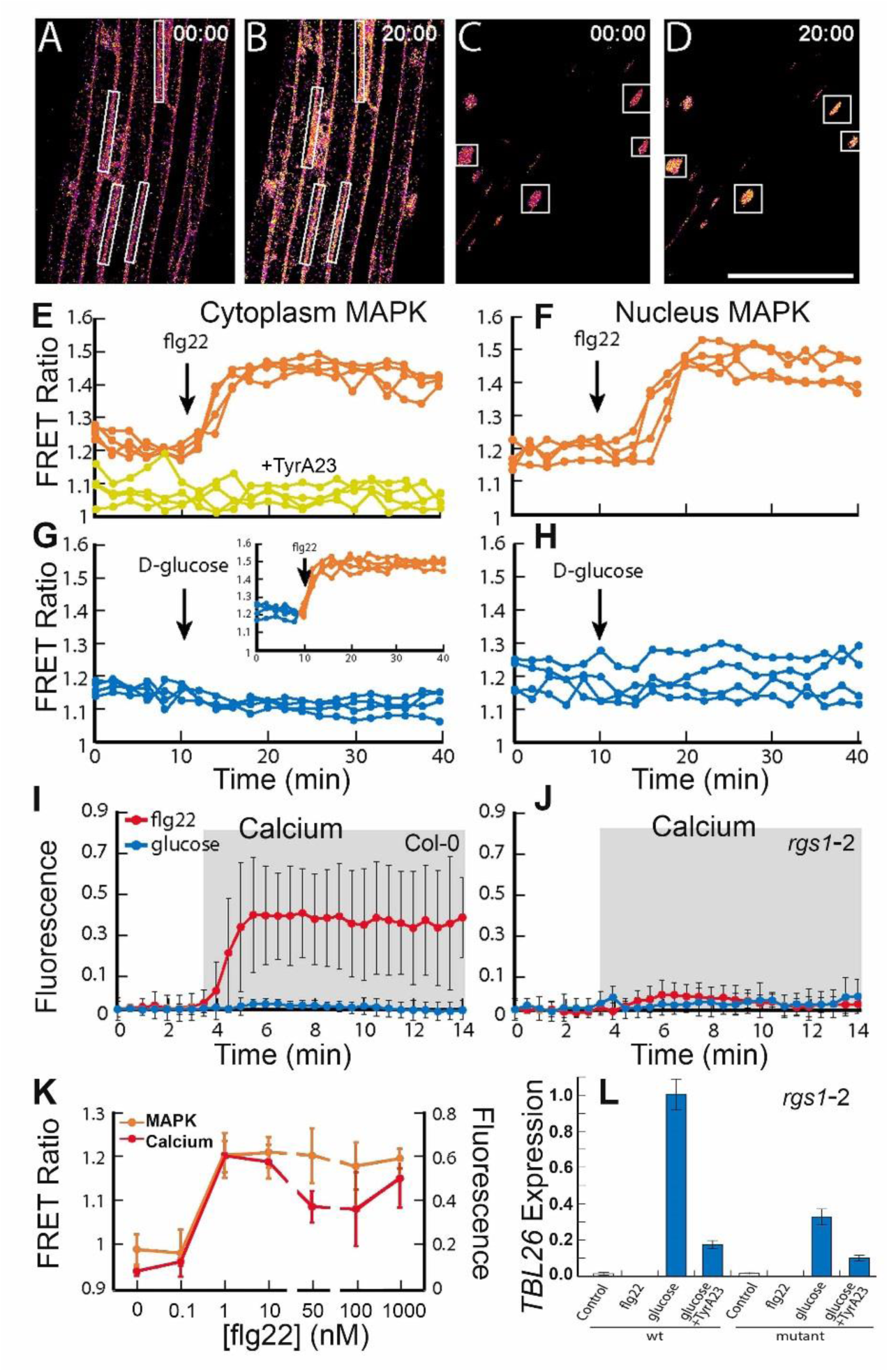
MAPK activation and Ca^2+^ signaling in response to flg22 vs. gene expression in response to D-glucose. (**A-D**) Processed confocal images of the epidermis of etiolated hypocotyls from the SOMA-NLS (**A, B**) and SOMA-NES (**C, D**) transgenic lines depicting the ratio of YPet to Turquoise GL emission produced by exciting Turquoise GL. Scale bar represents 100 µm. Time stamps indicate when the image was collected labeled minutes:seconds. Images at 00:00 were collected before treatment, while those at time point 20:00 were collected 5 minutes after treatment with 1 µM flg22. White rectangle represent regions of interest (ROIs) used to measure YPet and Turquoise GL emission. (**E-H**). The ratio of YPet to Turquoise GL emission produced by exciting Turquoise GL over time was determined using the ROIs shown in (**A**-**D**). During the first 10 minutes of each experiment the samples were incubated in water. The arrow indicates the time at which 1 µM flg22 (**E, G**) or 6% D-glucose (**F, H**) was added to the sample. (**E**) SOMA-NES lines pretreated with water (orange) or 50 µM TyrA23 (yellow) prior to imaging. Inset graph (**G**) shows SOMA-NES line pretreated with 6% D-glucose for 30 minutes prior to imaging. (**I**) flg22 dose-response in SOMA-NES lines. flg22 was added after two minutes of imaging. (**J, K**) R-GECO1 lines treated with flg22 or D-glucose in Col-0 (**J**) or *rgs1-2* (**K**) backgrounds. Fluorescence intensity changes of R-GECO1 in ∼20 regions of interests in wild type plants. Fractional fluorescence changes (ΔF/F) for R-GECO1 were calculated from background corrected intensity values of R-GECO1 as (F − F0)/F0, where F0 represents the average fluorescence intensity of the baseline of a measurement of each genotype. Error bars are standard deviations. Asterisks represent statistical significance (*P* < 0.01) between treatment and water as determined by 2-way ANOVA. (**K**) flg22 dose response in R-GECO1 and SOMA-NES lines after treatment for two or ten minutes respectively. (**L**) Quantification of *TBL26* gene expression by RT-qPCR in Col-0 and *rgs1-2* after treatment with water, flg22, or D-glucose. TyrA23 inhibitor treatment occurred for 1 hour prior to the ligand treatment.

We tested the role of D-glucose and flg22 in triggering an increased Ca^2+^ transient response using the intensity-based Ca^2+^ sensor R-GECO1, a red-shifted intensity-based Ca^2+^ reporter ^74^. We used stable transgenic Arabidopsis lines in wild type and *rgs1-2* backgrounds expressing cytosolic- and nuclear-localized R-GECO1. For our assay, D-glucose and flg22-induced Ca^2+^ changes in wild type and *rgs1-2* plants were measured over a time course in etiolated hypocotyls and normalized against the untreated samples (Fig 2I and 2J). Ca^2+^ levels represented by fractional fluorescence changes (ΔF/F; the difference between the fluorescence intensity before and after flg22 application divided by initial fluorescence intensity) increased in wild type (p< 0.01) while ΔF/F was greatly diminished (ΔF/F = ∼0.01) in the *rgs1*-2 mutant in response to flg22. D-glucose treatments did not significantly alter the ΔF/F in Col-0 wild type or *rgs1-*2 mutant (p=0.62), suggesting that D-glucose does not utilize AtRGS1 in a Ca^2+^ branch of the pathway. Activation of MAPK and the Ca^2+^ increase both occurred at a threshold of 1 nM flg22 (Figure 2K). Despite both D-glucose and flg22 inducing AtRGS1-internalization, these results show that within this context, only flg22 is capable of inducing MPK6 activity and Ca^2+^ changes.

Here, we showed that D-glucose signaling mediated by AtRGS1 does not involve MAPK or Ca^2+^ branches, however, D-glucose elicits a change in expression of a small set of genes over time and dose ^29,41^. *TRICHOMELESS LIKE 26* (*TBL26*), which is induced by D-glucose in an AtRGS1/G protein complex-dependent manner ^41^, is used as a reporter for activation ^2,29,75^. However, what is not known is if flg22 operates on this transcriptional pathway. As shown in Figure 2L, flg22 does not induce *TBL26* expression indicating separate outputs for D-glucose and flg22.

Paradoxically, the loss of AtRGS1 inhibits *TBL26* expression indicating that genetically, AtRGS1 acts like a positive modulator of signaling which is contradictory to our understanding that RGS proteins negatively modulate G protein activation. One solution to this paradox is that AtRGS1 has a positive regulatory role at the endosome; specifically that AtRGS1 endocytosis is essential. To test this, we quantitated glucose-induced *TBL26* expression in the presence of an endocytosis inhibitor Tyrphostin A23 (**TyrA23**) and found that *TBL26* expression is dramatically reduced (Figure 2L). This is consistent with the notion that AtRGS1 signaling has an endosomal origin, analogous to some GPCRs ^76^.

### AtRGS1 endocytic pathway is signal biased

Having shown that extracellular flg22 is detected by BAK1/FLS2 ^32,55,77^ and induces AtRGS1 endocytosis ^32^ and that extracellular D-glucose acts upstream of AtRGS1 endocytosis, both requiring at least in part the phosphorylation of AtRGS1 at its C terminal tail ^29,32^, we turned to the question of whether these two signals activate the same physical pool of AtRGS1. Endocytosis in plants utilizes two primary pathways: CME and SDE. Both pathways have been individually associated with the activity of specific proteins and responses ^49,78–80^; for example, CME and SDE cooperatively regulate the activity of RbohD in the flg22 pathway ^50^. We used high-resolution microscopies to achieve the needed spatial and temporal resolution to answer quantitative differences in signal-induced activation. To determine if one or both pathways regulate AtRGS1 activity at the membrane, we induced internalization with D-glucose and *flg22* while simultaneously inhibiting the CME pathway by arresting AP2µ function with TyrA23 ^79,81^ or inhibiting the SDE pathway by suppressing microdomain formation via sterol solubilization with methyl-β-cyclodextrin (**MβCD**) ^82,83^. MβCD, even at great excess, did not block *flg22*–induced AtRGS1 internalization compared to the control (p<0.01) (Figure 3A). Conversely, AtRGS1 internalization induced by glucose was reduced nearly 50% with a minimum concentration of 5mM MβCD (Figure 3B). This suggests that roughly half of the D-glucose-regulated pool of AtRGS1 is located in a SDE domain. To determine if the CME pathway regulates AtRGS1 activity, we applied TyrA23 with *flg22* or D-glucose. AtRGS1 internalization with both activators was significantly reduced (p<0.01); completely for flg22-induced AtRGS1 endocytosis (Figure 3C) and by 50% for D-glucose-induced AtRGS1 endocytosis (Figure 2D). The structurally similar but inactive TyrA23 analog TyrphostinA51 showed no significant effect on AtRGS1 internalization (p<0.01) (Figure 3C and 3D, TyrA51), indicating that the inhibitory effect of TyrA23 is chemically specific ^81^. When both inhibitors were applied simultaneously with glucose, AtRGS1 internalization was ablated (Figure 3E). The observation that TyrA23 decreased D-glucose-induced AtRGS1 endocytosis by only half compared to complete inhibition for flg22-induced endocytosis suggests that TyrA23 is not inhibiting through global cytotoxicity.

**Figure 3.**
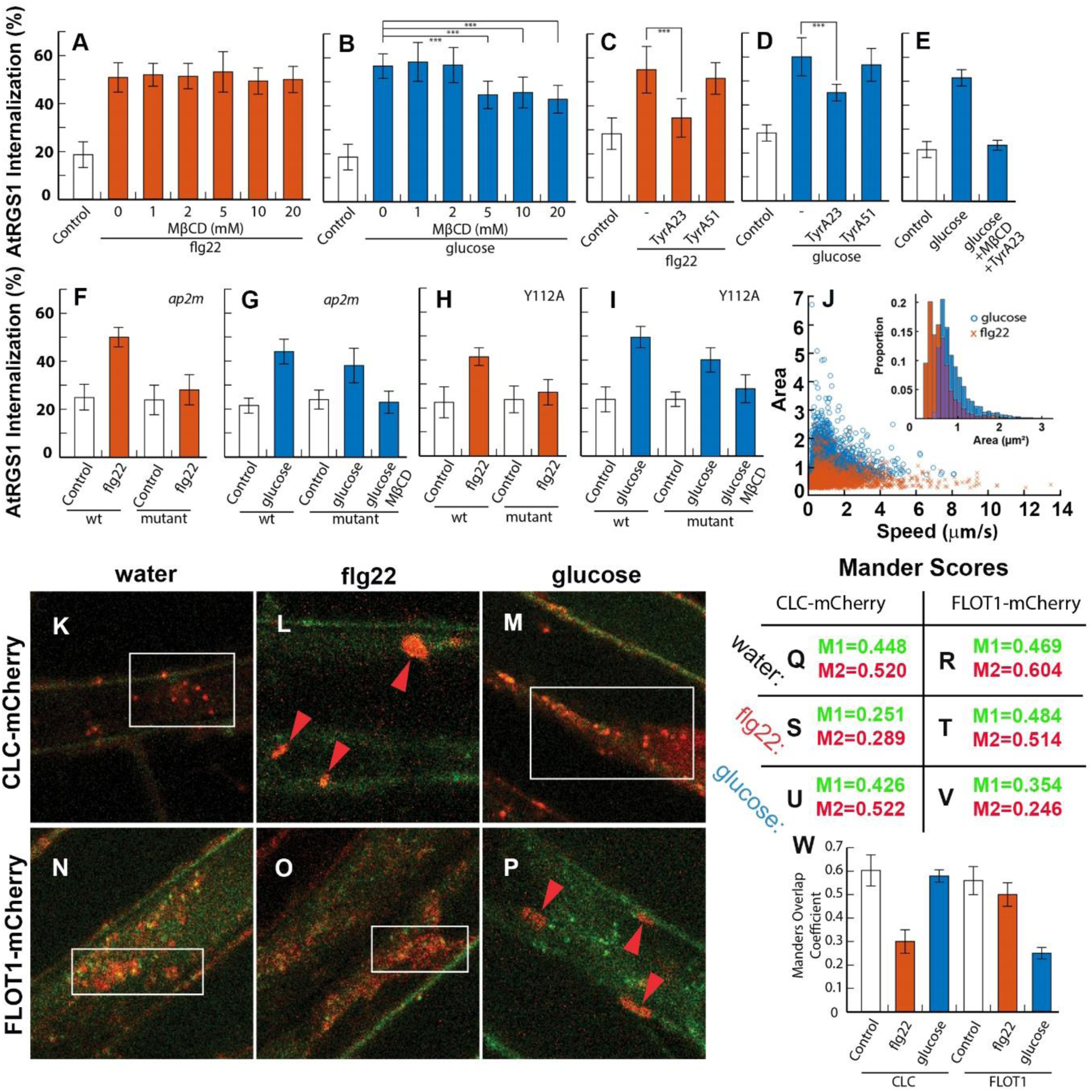
Two origins of AtRGS1 endocytosis. Pharmacological inhibitors show two origins of AtRGS1-YFP endocytosis. (**A**) AtRGS1-YFP seedlings were treated with increasing concentrations of MβCD at 0 mM (n= 18 (flg22) and 15 (glucose)), 1 mM (n= 14 and 14), 2 mM (n= 15 and 16), 5 mM (n= 17 and 14), 10 mM (n= 12 and 17), and 20 mM (n= 15 and 6) for 45 min followed by incubation in the same solution but supplemented with 1 μM flg22 for 10 min or (**B**) 6% D-glucose for 30 min before imaging epidermal cells. Internalized AtRGS1-YFP was quantified to determine total endocytosis of AtRGS1. MβCD does not inhibit flg22-induced AtRGS1 endocytosis, but does partially inhibit glucose-induced AtRGS1 endocytosis at 5mM and above (p<0.01). (**C**) Tyrphostin A23, an inhibitor of CME, significantly impairs flg22-induced AtRGS1 endocytosis (n=14) and (**D**) partially inhibits glucose-mediated AtRGS1 endocytosis (n=27) (p<0.01). The structurally similar, but inactive analog of TyrA23, TyrA51, has no effect (n=19) indicating that the effect of TyrA23 is specific. (**E**) When both inhibitors are applied with glucose, AtRGS1 internalization is reduced to basal levels (n=8) (p<0.01). A genetics approach to inhibit AtRGS1 internalization confirms pharmacological results. (**F**) A genetic null mutant of *ap2µ*, the cargo recognition complex for CME, results in complete inhibition of flg22-induced AtRGS1 (n=16) (p<0.01) down to basal levels, confirming TyrA23 results. (**G**) Glucose-induced internalization of AtRGS1 is partially inhibited in the *ap2µ* mutant (n=32) (p<0.01), but further reduced to basal levels with the addition of MβCD (n=18) (p<0.01). (**H**) Mutation of Y^112^ to Alanine in the AtRGS1 tyrosine motif recognized by ap2µ, AtRGS1^Y112A^, inhibits flg22-induced AtRGS1 endocytosis to basal levels (n=16) (p<0.01). (**I**) Glucose-mediated AtRGS1 endocytosis is partially inhibited in AtRGS1^Y112A^ (n=12) (p<0.05), but subsequently reduced to basal levels with the addition of MβCD (n=9) (p<0.01). Quantification of AtRGS1-YFP fluorescence is discussed in methods. (**J**) The speed and surface area of AtRGS1-GFP particles as tracked and measured by IMARIS from 30 second time lapse imaging using TIRF at 5 minutes after treatment with glucose and flg22. Identifiable AtRGS1-GFP particles are significantly smaller in the flg22 treated population (n=2026) compared to glucose (n=4619). No significant difference in speed is observed between particles in the two treatments. (**K-P**) Endocytosis markers CLC-mCherry and FLOT1-mCherry localize to the cell periphery and increase overlap with RGS1-GFP in a ligand dependent manner after treatment with flg22 and glucose. Zeiss confocal micrographs show AtRGS1-GFP (**green channel**) and either CLC-mCherry (**red channel K-M**) or FLOT1-mCherry (**red channel N-P**) after 5 minute treatments with water (**K and N**), flg22 (**L and O**), and glucose (**M and P**). After water treatment, CLC-mCherry (**K box inset**) and FLOT1-mCherry (**N box inset**) remain distributed throughout the cell cytoplasm and highly overlapped AtRGS1-GFP particles (**Q**). After treatment with flg22, CLC-mCherry migrates to the cell periphery with an observed increase in protein agglomeration (**L red arrows**) and decrease overlap score with RGS1GFP (**S**), while FLOT1-mCherry has no observable change compared to water (**G box inset and R**). Upon treatment with glucose, FLOT1-mCherry protein bodies congregate at the cell periphery (**P red arrows**) and decrease overlap with RGS1-GFP (**V**), while CLC-mCherry remains unchanged compared to water (**M box inset and U**). (**W**) The average overlap scores for CLC-mCherry and FLOT1-mCherry after addition of water (n= 8 and 9), flg22 (n= 4 and 7), and glucose (n= 4 and 3). flg22 addition induces a significant change in CLC compared to glucose and water while glucose induces a significant change with FLOT1 compared to flg22 and water (p<0.01)

These results suggest that there is a single flg22-induced pool that internalizes through a CME pathway and that there are two distinct glucose-induced pools, one internalizes through the CME pathway and the other through the SDE pathway. Because the glucose-induced pool is equally inhibited by the CME and SDE inhibitors, a rapid equilibrium between the pools is not likely, otherwise neither inhibitor would have shown efficacy.

The use of TyrA23 as a specific endocytosis inhibitor is controversial ^84–86^, therefore we added a genetic approach by measuring AtRGS1 internalization in the AP-2µ null mutant, *ap2m* ^87^. The AP-2 complex plays a critical role in transporting cargo in the CME pathway, whereby the µ subunit of this tetrameric AP-2 complex directly interacts with cargo proteins during endocytosis ^45,88^. In the *ap2m* mutant seedlings, flg22-induced AtRGS1 internalization was ablated (p<0.01), consistent with the TyrA23 results (Figure 3F). In contrast, D-glucose induced about half AtRGS1 internalization in the *ap2m* mutant compared to Col-0 wild type (p<0.01) also consistent with the TyrA23 result. The addition of MβCD with D-glucose ablated AtRGS1 internalization in the *ap2m* mutant to basal levels (Figure 3G). These observations suggests that there are two signal-dependent pools of AtRGS1 on the plasma membrane; 1) a homogenous pool for flg22 signaling and 2) a conglomerate pool for D-glucose signaling.

Because D-glucose utilizes both CME and sterol-dependent pathways to induce AtRGS1 internalization, we tested if the depletion of CME pools of AtRGS1 by D-glucose would alter flg22-induced MPK6 activation. After pretreating SOMA-NES lines with D-glucose for 30 minutes prior to imaging, we found that D-glucose did not have an effect on flg22-increased FRET efficiency (Fig 2G inset) suggesting that the glucose pool of AtRGS1 is sequestered from the flg22-induced CME of AtRGS1.

To determine if CME-mediated endocytosis regulates flg22-induced MPK6 activity, we pretreated SOMA-NES lines with TyrA23. TyrA23 successfully blocked the increases in FRET efficiencies that were observed in the absence of the inhibitor (Fig 2E, replicates in Fig S2 G-N). The negative control TyrA51 showed no significant effect on FRET efficiency after treatment with flg22 (Fig S2 P-V), indicating that the inhibitory effect of TyrA23 is chemically-specific to its role in blocking CME-mediated endocytosis and suggesting that AtRGS1 endocytosis, per se, is required. However, this observation does not exclude the possibility that within the context of flg22-induced MAPK signaling that some other endocytosed protein is solely rate-limiting for MAPK cascade activation. This possibility is excluded in the following section.

### A point mutation in the intracellular loop disarms biased signaling

The µ subunit of the AP-2 complex is a cargo recognition and interaction component in the CME pathway. It binds to known tyrosine motifs, YXXΦ, where X is any amino acid and Φ is an amino acid with a hydrophobic side chain ^89^. The second intracellular loop in AtRGS1 contains such a motif with the amino acid sequence Y^112^FIF ^90^. To determine if this motif is necessary for AtRGS1 endocytosis and a potential binding motif for AP-2µ, we generated AtRGS1 with a tyrosine to alanine mutation (AtRGS1^Y112A^). flg22 failed to induce endocytosis of the AtRGS1^Y112A^ mutant (Figure 3H). D-glucose-induced internalization of the AtRGS1^Y112A^ was reduced to half (p<0.05) compared to wild type internalization and was further reduced to the resting level with the addition of MβCD (p<0.01) (Figure 3I) demonstrating that the Y^112^FIF is necessary for internalization and likely a recognition and binding site for AP-2µ. The flg22-induced AtRGS1 pool is entirely mediated by the CME pathway whereas roughly half of the D-glucose-induced AtRGS1 pool internalizes through the CME pathway. While we recognize that both TyrA23 and genetic ablation of AP-2µ have off-target effects, the point mutation in the AtRGS1 cargo recognition motif provides strong evidence for flg22- and glucose-inducing CME of distinct pools of AtRGS1.

### Physically-distinct, dynamic populations of AtRGS1: Architectural basis for biased signaling

We showed through pharmacological and genetic approaches that two signal-dependent pools of AtRGS1 exist, raising the possibility that the two AtRGS1 pools are physically distinct on the cell membrane. The different dependency on sterol and clathrin for partial D-glucose- vs. flg22-induced endocytosis of AtRGS1 suggests that this is the case. To test this hypothesis, we imaged AtRGS1 with a C-terminal GFP tag using Total Internal Reflection Microscopy (**TIRF**) and IMARIS (v9.2.2, Bitplane Inc) surface tracking (Figure S3 A-E, examples of raw data). We used TIRF because it provides excellent signal to noise ratio for visualizing only fluorescent proteins near and on the membrane by illuminating with an evanescent wave of excitation light that quickly decays beyond the cell membrane. While we cannot distinguish membrane bound AtRGS1-GFP from AtRGS1-GFP located at the extreme membrane periphery, we can quantitate the intrinsic properties of AtRGS1-GFP populations under different ligand conditions. We calculated the average size and speed of GFP fluorescence during a 30-second interval taken from time-lapse imaging at 5 and at 15 minutes post treatment with glucose and flg22. Two clearly distinct, signal-dependent populations of AtRGS1 were observed (Figure 3J). After 5 minutes under glucose conditions, the surface area of AtRGS1-GFP clusters were significantly larger (ẍ = 0.9403 µm^2^ n=4619) than flg22-treated cells (ẍ = 0.5998 µm^2^ n=2026) (Figure 3J inset) (p<0.01). At 15 minutes, the area of these flg22-treated clusters increased slightly to 0.6733 µm^2^ (n=1751), and the area of the glucose-treated clusters increased to 1.0072 µm^2^ (n=6209) (Figure S3 F-H). For flg22-treated cells, the average velocity of the clusters were 1.53µm/s and 1.63µm/s at 5 and 15 minutes, respectively (n= 2026 and 1751) and for the flg22-treated cells averages, the velocities were 0.98µm/s and 1.01µm/s at 5 and 15 minutes, respectively (n= 4619 and 6209). Two populations of differently-sized AtRGS1 protein clusters moving at different speeds provides physical evidence supporting two origins of endocytosis.

### The signal dependency correlates with endocytosis marker redistribution

The Clathrin Light Chain (**CLC**) and Flot1 proteins are associated with CME and SDE, respectively ^42,49^. We generated transgenic lines expressing AtRGS1-GFP with either CLC-mCherry (CME) or Flot1-mCherry (SDE) endocytosis markers to investigate the localization of AtRGS1 in relation to both markers. We imaged the response of both endocytosis markers 5 minutes after application of water, flg22, and glucose to look for changes in marker distribution in the cell (Figure 3K-P). Using Manders Overlap Coefficient (hereafter, Manders), we quantified the co-occurrence of both endocytosis markers with AtRGS1-GFP under all treatment conditions. We used Manders instead of Pearson’s Correlation because qualitative analysis showed a ligand-dependent change in marker localization and shape. Manders provides a quantitative measure of the change in overlap of the two signals, not the change in signal intensity (Pearson’s Coefficient) that may be a result of AtRGS1 internalization (e.g. change in compartment pH) and not correlated to direct interaction with either endocytosis marker.

A subset of the Manders after background subtraction are shown in Figure 3Q-V and correspond to the confocal micrographs in Figure 3K-P. M1 represents the proportion of total GFP from AtRGS1 that overlaps with mCherry. Similarly, M2 represents the proportion of total mCherry from CLC or FLOT1 that overlaps with GFP. Under conditions without either flg22 or D-glucose (Figure 2K and 2N), AtRGS1-GFP and our endocytosis markers have a high overlap baseline (Figure 3Q and 3R). After flg22 addition (Figure 3L and 3O), the proportion of CLC associated with AtRGS1 decreased, with the proportion of AtRGS1 overlapping with CLC also decreasing significantly (p<0.01). Simply stated, a smaller defined subset of the AtRGS1 and CLC populations are overlapping with each other in a ligand-dependent manner with the endocytosis marker also exhibiting a structural change. This response was not observed with FLOT1 when flg22 was added (Figure 3T). With glucose addition (Figure 3M and 3P), CLC showed no significant change compared to water (Figure 3U), while FLOT1 overlap with RGS1 decreased compared to the water baseline (p<0.01) (Figure 3V). The same responses were observed at 15 minutes post ligand addition (Figure S3 I-T). The averages for the entire collection of M2 scores from all samples are presented in Figure 3W. These results suggests that AtRGS1 association with the CME and SDE-defined routes is biased toward the activating signal. In summary, the different intrinsic trafficking properties further support at least two pools of ligand-activated AtRGS1.

### Signaling bias involves phosphorylation from different kinases

Phosphorylation of AtRGS1 is a necessary requisite for both D-glucose- and flg22-induced endocytosis. The C-terminus of AtRGS1 contains multiple di-serine residues that could function as potential phosphorylation sites for signal transduction. A truncated version of AtRGS1 lacking the 43 C-terminal residues, AtRGS1^ΔCt^, served as a blunt instrument for a phosphorylation-deficient mutation to determine if the C-terminal tail is necessary for AtRGS1 endocytosis (Figure S4A). Application of flg22 failed to internalize the AtRGS1^ΔCt^ mutant compared to water control (p<0.01) (Figure 4A). D-Glucose application however, internalized the AtRGS1^ΔCt^ mutant, but at 50% the level of the wild type AtRGS1 (p<0.01). MβCD completely inhibited internalization of the AtRGS1^ΔCt^ mutant levels (p<0.01) (Figure 4B). These results show that the C-terminus is necessary for flg22- and glucose-induced endocytosis of AtRGS1 and support two subpopulations of AtRGS1 among the glucose-induced pool.

**Figure 4.**
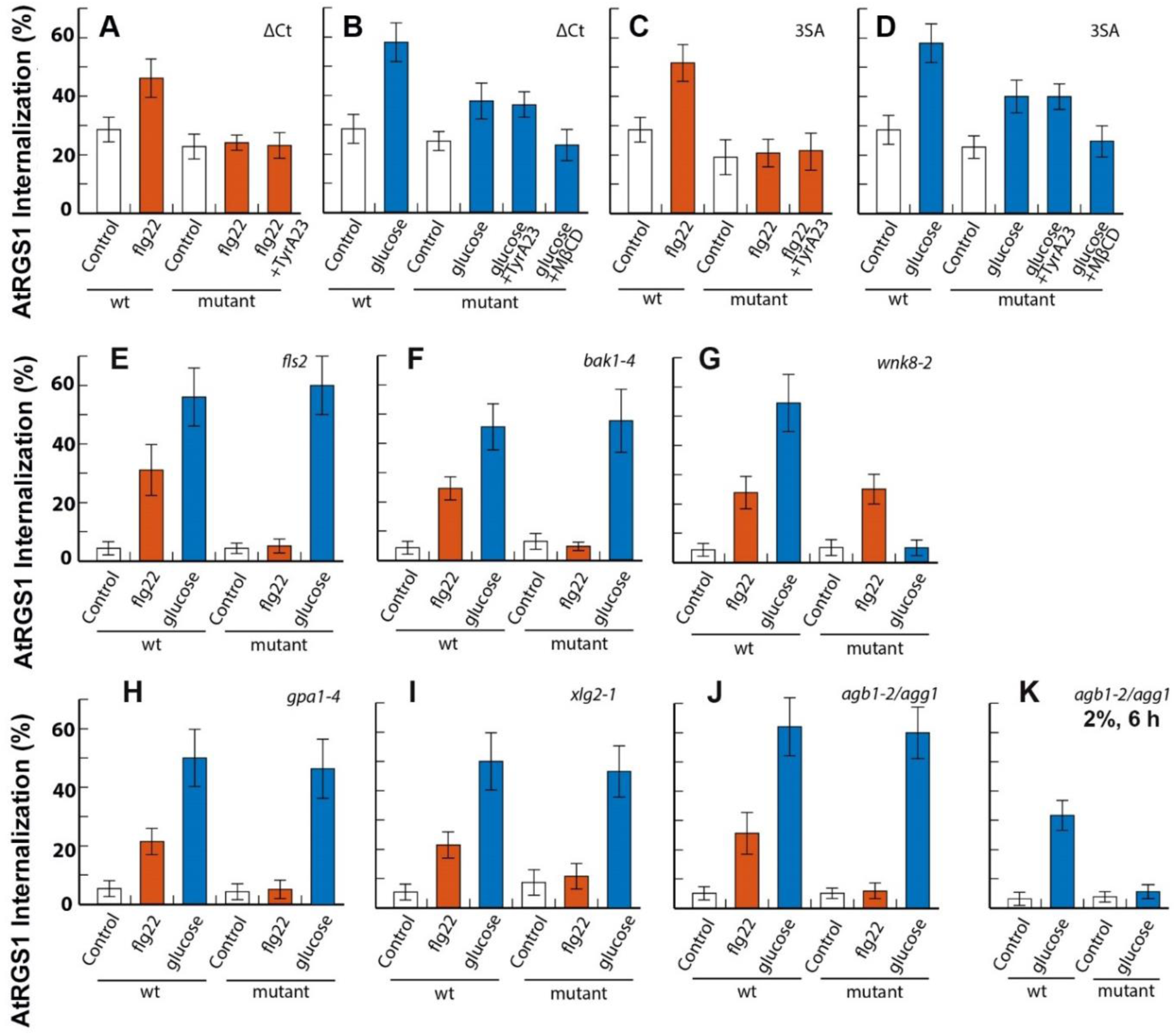
CME of AtRGS1 is phosphorylation dependent and G protein specific. Known phosphorylation sites at serine residues on the C-terminal end of AtRGS1 are required for both flg22 and glucose-induced CME of AtRGS1. (**A**) A truncated AtRGS1, AtRGS1^ΔCt^, lacking the 43 most C-terminal residues, including 8 serines, resulted in basal levels of flg22-induced AtRGS1 internalization (n=11) (p<0.01). (**B**) Glucose-induced AtRGS1 internalization is partially reduced in AtRGS1^ΔCt^ (n=12) (p<0.01), but further reduced to basal levels with the addition of MβCD (n=13) (p<0.01). (**C and D**) Mutation of three specific serine residues to alanine at 428, 435, and 436, in a full length AtRGS1, AtRGS1^3SA^, yielded similar results to AtRGS1^ΔCt^ for flg22 (n=10) and glucose-mediated internalization (n=22) (p<0.01), but indicate phosphorylation of one or several specific serine residues is necessary for CME of AtRGS1. flg22 and glucose require known kinases for AtRGS1 internalization. (**E**) Genetic ablation of the flg22 receptor FLS2 yields basal levels of flg22-induced AtRGS1 internalization (n=12) (p<0.01), but does not affect glucose-induced AtRGS1 internalization (n=9). (**F**) Similarly, null mutation of the BAK1 co-receptor, *bak1-4*, results in ablated flg22-induced AtRGS1 internalization (n=16) (p<0.01), but not for glucose (n=17) (p=0.36). (**G**) The high dose and low duration glucose specific WNK8 kinase is necessary for glucose induced AtRGS1 internalization (n=13) (p<0.01), but not flg22 (n=10) (p=0.9). Individual G proteins are necessary for AtRGS1 endocytosis in a ligand-specific manner. (**H**) A genetic null mutant of Gα, *gpa1-4*, limits flg22-induced endocytosis of AtRGS1 to basal levels (n=15), while glucose-induced endocytosis is unaffected compared to wild-type (n=22). (**I**) A null mutation of XLG2, *xlg2-1*, an extra-large Gα protein, also reduced flg22-induced AtRGS1 endocytosis to basal levels (n=14), but had no significant effect on glucose-mediated endocytosis compared to wild type (n=19) (p<0.01). (**J**) A null mutation of the AGB1/AGG1 heterodimer, *agb1-2/agg1*, inhibited flg22-induced AtRGS1 endocytosis (n=15), but had no effect on high dose, low duration glucose-induced AtRGS1 endocytosis (n=17) (p<0.01). (**K**) At low dose, long duration AGB1/AGG1 are necessary for glucose induced internalization (n=25). Quantification of AtRGS1-YFP fluorescence is discussed in methods.

Two phosphorylated serine residues on the C-terminus of AtRGS1 at positions 428 and 435 or 436 are necessary for at least partial endocytosis of AtRGS1 induced by both D-glucose and flg22. Therefore, we mutated these serines to alanine (AtRGS1^3SA^) to determine if these specific residues were necessary to induce internalization by either agonist. Upon treatment with flg22, AtRGS1^3SA^ internalization was at basal levels compared to wild type (p<0.01) (Figure 4C), confirming previous results by Tunc-Ozdemir *et al* ^32^ and suggesting that the CME pathway is dependent on phosphorylation of serines 428, 435 and/or 436. In the case of glucose, endocytosis was only partially impaired by the AtRGS1^3SA^ mutant (p<0.01) (Figure 4D). MβCD completely blocked glucose-induced endocytosis of the AtRGS1^3SA^ mutant (Figure 4D), consistent with our previous results that show glucose-induced internalization utilizes both CME and SDE pathways and that the CME pathway requires phosphorylation at Ser_428/435/436_.

As an added note, these various genetic results enable us to exclude an osmotic effect that effectively sequesters a portion of the plasma membrane pool of AtRGS1 into a sterol-dependent microdomain that is incapable of internalizing its components.

### System bias by skewed kinase and G-protein complex composition

Because the cluster of phosphorylated serines on the C-terminus of AtRGS1 are required for AtRGS1 internalization, we hypothesized that one mechanism for system bias is to functionally-sequester cognate kinases for D-glucose and flg22 into their respective ligand-delineated pools. To test this hypothesis, we quantified AtRGS1 internalization in the mutant backgrounds of the flg22-activated FLS2 kinase (*fls2*) and BAK1 co-receptor (*bak1*) and the D-glucose-activated WNK kinase (*wnk8-*2, *wnk1*-1) null mutants. Upon treatment with flg22, AtRGS1 endocytosis was ablated in *fls2* and *bak1-4* mutants (P<0.01), while D-glucose-induced internalization in these mutants was unaffected (P=0.70 and 0.37 respectively) (Figure 4E and 4F). In contrast, D-glucose-induced internalization in the *wnk8-2* was ablated (p<0.01), while flg22-induced AtRGS1 endocytosis was unaffected (P=0.89) (Figure 4G). Our results suggest that FLS2 and BAK1 are specific to the CME pathway and WNK8 kinase is specific to the SDE pathway.

The canonical components of the heterotrimeric G protein complex are necessary for D-glucose-induced internalization of AtRGS1 ^2,29^. More specifically, Gβγ is required for the recruitment of WNK kinases for phosphorylation of AtRGS1, leading to AtRGS1 endocytosis and activation of downstream G signaling. We hypothesized that individual components of the G protein heterotrimer may be required for biased signaling. To test this hypothesis, we quantified AtRGS1 endocytosis in G protein mutant backgrounds after flg22 and glucose activation. In the AtGPA1 (Gα) null background, *gpa1-4,* AtRGS1 endocytosis showed no significant difference compared to wild type when glucose was applied (p=0.48) (Figure 4H). After addition of flg22, however, AtRGS1 endocytosis was at basal levels (p < 0.01) in the absence of Gα indicating that the G subunit is essential for this pathway (Figure 4H). We additionally tested AtRGS1 in the absence of XLG2, one member of a three-gene family of atypical G proteins. XLG proteins have a homologous C-terminal Gα domain and an N-terminal nuclear localization signal ^91^. Loss of XLG2 in the *xlg2-1* mutant did not affect AtRGS1 endocytosis by D-glucose application (p=0.43), but completely inhibited AtRGS1 endocytosis to basal levels after addition of flg22 (p<0.01) (Figure 4I). In the *agb1-2/agg1* double null mutant, AtRGS1 endocytosis was diminished after addition of flg22 (p<0.01) (Figure 4J). Loss of the Gβγ dimer in the *agb1-2/agg1* mutant had no effect on glucose-induced AtRGS1 endocytosis compared to wild type after 30 minutes of treatment with 6% D-glucose (p=0.79) (Figure 4J). Interestingly, activation by a lower dose (2%) of D-glucose over a longer duration (6 hour) required AGB1/AGG1 (p<0.01) (Figure 4K).

### VPS26: a candidate plant β-arrestin-like adaptor necessary for AtRGS1 endocytosis in the CME pathway

Higher plant genomes do not encode canonical (i.e. visual and nonvisual) arrestins ^92^. Therefore, we sought candidate adaptors for AtRGS1 that may function like β-arrestins to recruit AP2/clathrin to AtRGS1 for endocytosis by querying 3-D structure databases. As shown in Figure S5A and B, we identified three *Arabidopsis* VACUOLAR PROTEIN SORTING 26 (VPS26) proteins that contain arrestin folds ^93^ and are orthologous to human VPS26. *Arabidopsis* VPS26A and VPS26B amino acid sequences are 91% identical whereas VPS26-like is 20% identical to VPS26A and VPS26B. In mammals and plants, VPS26 operates with VPS29 and VPS35 in the retromer complex on the endosome ^94^. This raises the possibility that VPS26 proteins have a moonlighting function in modulating AtRGS1 internalization.

To compare the arrestin and AtVPS26A protein structures, we first created a model of AtVPS26A using MODELLER ^95^. Toward this, a high quality homogeneous sequence alignment was generated using the VPS26 family (Arabidopsis VPS26A, VPS26B, VPS26like and *Homo sapiens* VPS26A) and arrestin family (Vertebrates: Human arrestin-1 and arrestin-2; bovine arrestin-1, arrestin-2, arrestin-2, and a variant p44; Squid arrestin-1; and the invertebrate shrimp arrestin (Figure S4A). Human, bovine, and squid sequences were included because PDB structures are available. The squid and shrimp sequences were added for divergence information (among the opistokonts). AtVPS26A and human VPS26A share 56.48% sequence identity while AtVPS26A and arrestins share only 14-17% sequence identities (Figure S4B). These results rationalized our use of the high-resolution (2.1Å) crystal structure of *Homo sapiens* Vps26a (PDB [2FAU]) rather than an arrestin structure to generate models of AtVPS26A for subsequent comparison to arrestin. Details of the top five selected models (atvps26a1 →5) are provided in the Methods.

We compared the atvps26a-2 model with the bovine arrestin-3 (PDB [3P2D ^96^]). Although the primary amino acid identity between arrestin and AtVPS26A is only ∼15% (Figure 5A), the structure of AtVPS26A model shows a remarkably similar arrestin scaffold comprised of a semi-symmetric fold of two β strand sandwich structures in the N domain and C domains linked by central loops with each sandwich formed by 3 or 4 β strands, respectively.

**Figure 5.**
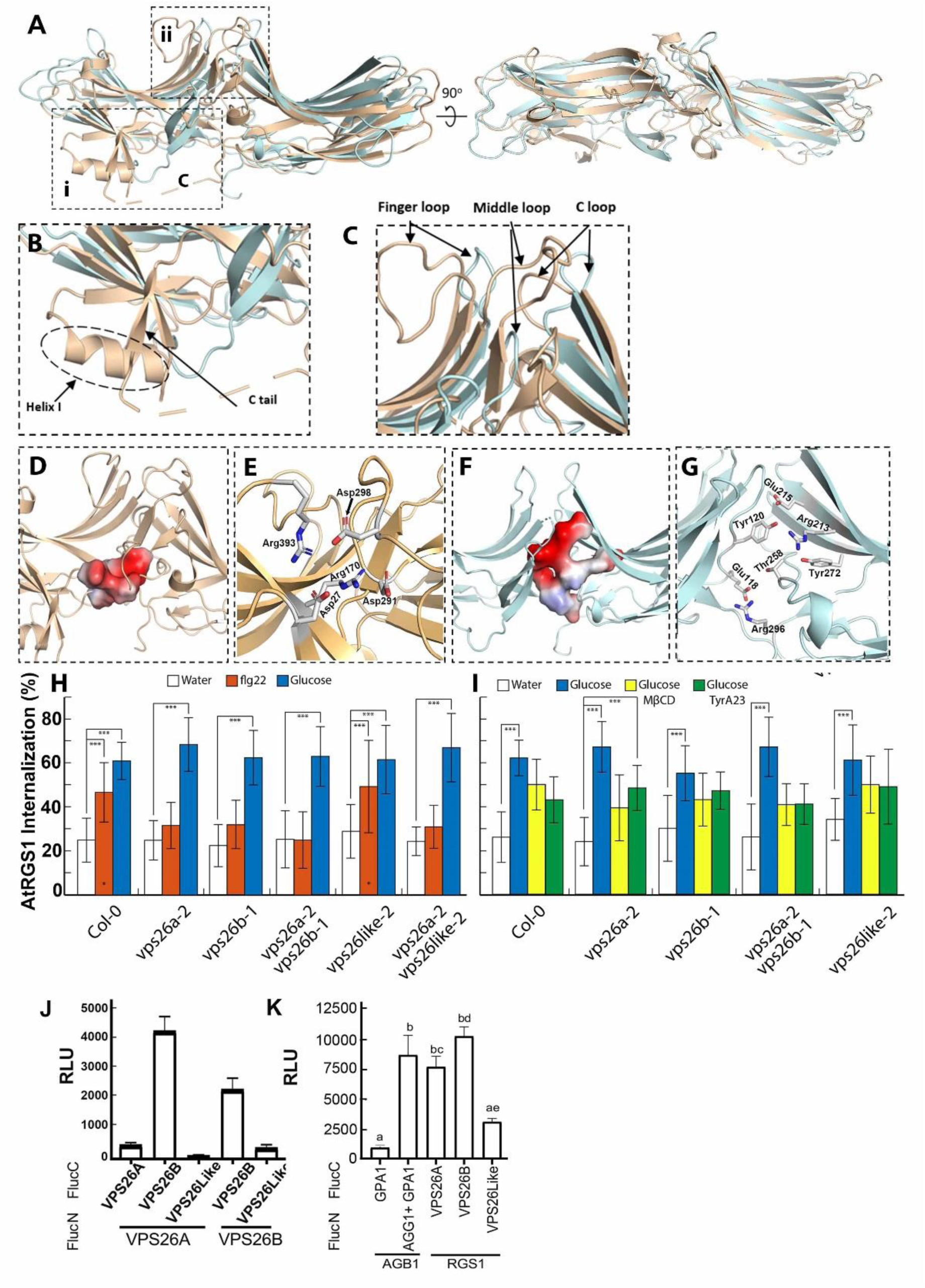
Arabidopsis VPS26 subunits of the retromer complex may moonlight as arrestin-like adaptors for AtRGS1 internalization. (**A)** 3-D alignment of the model atvps26a-2 (colored cyan) with the bovine arrestin-3 (PDB 3P2D, (colored wheat). The structure of model atvps26a-2 shows a similar arrestin scaffold with arrestin-3 which contains a semi-symmetric fold of two β strand sandwich structures in the N domain and C domains linked by the central loops (ii); each sandwich is formed by 3 or 4 β sheets, respectively. (**B)**. Differences between atvps26a-2 model and bovine arrestin-3. Model atvps26a-2 lacks a short α-helix (i) inside the arrestin N-terminal domain which has been implicated in receptor binding. Arrestin-3 contains a longer C terminal tail which extend all the way to bind the N terminal domain which is important for clathrin-mediated endocytosis (CME) in animals. The C terminus of atvps26a-2 has no extension. (**C**). The central crest. The central crest of atvps26a is similar with that of arrestin-3 which includes the finger loop, middle loop and the C loop, although arrestin-3 has a longer finger loop which is important for receptor binding. (**D, E**) The polar core of bovine arrestin-3.The residues in the polar core of arrestin-3 are Asp_27_, Arg_170_, Asp_291_, Asp_298_ and Arg_393_ which are shown as vacuum electrostatics (**D**) and sticks (**E**). (**F,G**) The polar core of AtVPS26A. The residues of atvps26a-2 are the N domain residues Glu_118_ and Tyr_120_, and C domain residues Arg_213_, Glu_215_, Thr_258_, Tyr_272_, and Arg_296_ which are also shown as vacuum electrostatics (**F**) and sticks (**G**). Although different amino acid composition, both cores consist of similarly-positioned, positive-charged residues and allow the formation of hydrogen bonds under physiological conditions. However, the orientation and shape of the polar core of atvps26a-2 is distinct from arrestin-3. The arrestin-3 polar core is embedded between β sheets in the N terminal domain whereas the atvps26a-2 core is open and elongated, spanning the length of space between the N domain and C domain. (**H**) Quantification of AtRGS1-YFP internalization in WT and *vps26* null mutants after treatment with water, 1 µM flg22 or D-glucose. (**I**) AtRGS1-YFP seedlings were pretreated with MβCD or TyrA23 prior to treatment with D-glucose. Error bars represent standard deviation. Data is averaged across three separate experiments. (**J**) Split luciferase complementation by VPS26A and VPS26B heterodimers and VPS26B homodimers. Error bars are standard error of the mean (SEM). Means with different letters indicate significantly different (Tukey’s HSD test, p<0.05). n=64 leaf discs from 4 individual tobacco plants. **K**) Split-luciferase complementation by VPS26A and AtRGS1. Positive control is complementation by the heterotrimeric G protein complex (AtGPA1/AGB1/AGG1). Negative control is AtGPA1 and AGB1 in the absence of AGG1. Error bars are SEM. n=64. (p<0.05).

For arrestin, the conserved structures mainly include N-terminal and C-terminal arms, a central crest which is comprised of a finger loop ^97^, a middle loop ^98,99^(see box ii of Fig. 5), and a C loop ^100^ (see box I of Fig. 5), a gate loop, a polar core, and the hinge domain ^101^. The N-terminal and C-terminal arms stabilize the arrestin conformation. Model atvps26a-2 shares similar N-terminal and C-terminal scaffolds with arrestin however it lacks a short α-helix inside the arrestin N-terminal domain (Figure 5B) which has been implicated in receptor binding ^102^. In addition, arrestin has a longer C-terminal tail, which extends to the N terminal domain, important for linkage needed to enable CME. The C terminus of atvps26a-2 has no extension; however, some arrestins also have a short C-tail ^103^.

The overall central crest of atvps26a-2 is similar to that of arrestin. It includes the finger loop, middle loop, and the C loop (Figure 5C). While arrestin has a longer finger loop important for receptor binding, this is intrinsically disordered as the rearrangement of the finger loop is a major change associated with arrestin activity ^104^.

The polar core is the key component of the phosphate sensor. In arrestin, the polar core is comprised of five charged side chains including two Arg and three Asp that are essential to its activation ^105^. The residues in the polar core of bovine arrestin-3 are: Asp_27_, Arg_170_, Asp_291_, Asp_298_ and Arg_393_ (Figure 5D and E). The human vps26A polar core residues are conserved N domain residues Glu_119_ and Tyr_121_, and C domain residues Lys_213_, Glu_215_, Thr_258_, Tyr_272_, and Arg_296_ ^106^. Based on this, we designated the polar core of atvps26a to be comprised of the N-domain residues Glu_118_ and Tyr_120_, and the C domain residues Arg_213_, Glu_215_, Thr_258_, Tyr_272_, and Arg_296_ (Figure 5 F and G). Although different in amino acid composition, both cores consist of positive charged residues and allow the formation of hydrogen bonds under physiological conditions. However, the orientation and shape of the polar core of VPS26 is distinct from arrestin. The arrestin polar core is embedded between β sheets in the N terminal domain whereas the VPS26 core is open and elongated, spanning the length of space between the N domain and C domain.

The most divergent structure from bovine arrestin-3 (3P2D) is a 3Ȧ-resolved truncated arrestin from squid (PDB 1CF1) ^107^. Unlike for bovine arrestin-3, the C-tail interaction with the gate loop of the polar core is short and flexible. This functional divergence between squid and human arrestin, both recruited to receptors, assuages the lack of a long C-tail in the atvps26a-2 model.

Taken together, these similarities between AtVPS26 and arrestin prompted the hypothesis that AtVPS26 is a candidate adaptor for CME of AtRGS1 or, in any case, necessary for the observed signaling bias. Therefore, we isolated null alleles of the three VPS26 genes in Arabidopsis: VPS26A, VPS26B, and VPS26-LIKE (aka VPS26C) ^94,108^ and phenotyped the hypocoty length at 5 days, the same age stage used for the previous assays (Figure 1-4). As shown in Figure S4F, there were no qualitative differences between VPS26 null mutants and wild type phenotype precluding any developmental basis for alterations in AtRGS1 activation in these mutants. As shown in Figure 5H, loss of either VPS26A or VPS26B dramatically reduced the flg22-induced internalization of AtRGS1 to levels that are not statistically different from the baseline level (p=0.34) whereas loss of VPS26-like had no statistical effect on AtRGS1 internalization by flg22 (p=0.65). Loss of any of the three VPS26 proteins had no statistical effect on D-glucose-induced internalization of AtRGS1 (p<0.001) suggesting that VPS26A/B are involved in flg22, but not glucose-induced AtRGS1 endocytosis. To confirm that VPS26 is not involved in D-glucose-induced AtRGS1 internalization, we quantitated the CME and SDE portions of this trafficking pathway in the *vps26* mutants. As shown in Figure 5I, in each of the *vps26* mutant backgrounds, both the TyrA23A and MβCD-inhibitable segments of the D-glucose-induced internalization of AtRGS1 remained intact. This suggests that the TyrA23-dileneated (CME) pathway used by AtRGS1 when activated by flg22 differs from the TyrA23A-dileneated pathway activated by D-glucose.

The genetic data that both the single null mutations of VPS26 A and B ablate flg22-induced AtRGS1 endocytosis suggest that VPS26A and B form obligate dimers. To test if VPS26A and B subunits heterodimerize *in vivo*, BiFC analysis was conducted and showed that heterodimers can form from ectopically-expressed monomers and suggested that the orientation is head-to-tail (Figure S5A). Self-association of arrestin family members also occurs and may be part of a regulatory mechanism for arrestin activation ^109^. For relative quantitation of *in vivo* interaction, split luciferase was performed and found that the strongest interaction was between VPS26A and VPS26B (Figure 5J). VPS26B dimers form under these conditions but not VPS26A dimers or any oligomer with VPS26Like subunits. We have yet to find conditions which allow stable expression of VPS26 subunits tagged with a full-length auto-fluorescent protein in Arabidopsis suggesting that the additional mass of the tag prevents dimer formation and that monomers are unstable.

To test if the full-length AtRGS1 protein interacts with the VPS26 subunits *in vivo*, BiFC, split luciferase, and Y2H analyses were performed. Fluorescent complementation of YFP by AtRGS1-nYFP with cYFP-VPS26A and with VPS26B-cYFP was observed (Figure S5B). The split luciferase assay confirmed that this interaction is as strong as the interaction between AtGPA1 and its AGB1/AGG1 partner (Figure 5K). Finally, Y2H showed that the interaction between AtRGS1 and VPS26 is direct. The entire C-terminal half of AtRGS1 (thus lacking the 7 TM domain) interacts with VPS26B (S5H). Surprisingly, removal of the post RGS-box C-terminal tail which includes the Ser_428,435,436_ phosphorylation cluster (Figure S5C) did not ablate this interaction however, additional loss of two other phosphorylated Ser located between helices VII and VIII of the RGS box completely abolished the interaction (Figure S5D).

## Discussion

The key finding in this study is system bias wherein a single receptor-like RGS protein modulates different quantifiable signaling outputs from two distinct signal inputs (Figure 6A). All of the measured signal outputs correspond to a specific ligand input. For example, flg22 induces Ca^2+^ release (Figure 2I-K) and MAPK signaling (Figure 2E-H, K) whereas D-glucose or its metabolite has a measured effect on gene expression (Figure 2L). For flg22, all members of the G protein heterotrimer, along with XLG2 and VPS26A/B are necessary for CME of AtRGS1. Additionally, specific phosphorylation of di-serines at the AtRGS1 C-terminus is required. What is interesting is that D-glucose signaling that transduces through the same CME of AtRGS1 also requires phosphorylation of the same di-serine residues, but not VPS26 for high dose and low duration sugar exposure (Figure 6B). This phenomenon of signaling through AtRGS1 mediated by the same mechanism of phosphorylation-dependent endocytosis with different G proteins and adapters raises the important question of how are extracellular ligands discriminated for downstream signaling events? Although AtRGS1 endocytosis is necessary for G protein activation it would seem that AtRGS1 is not the discriminator; rather an RLK protein that directly phosphorylates AtRGS1 ^30,32^, AtGPA1 ^110^, and potentially VPS26 provides the requisite information for ligand-specific downstream signal transduction in the cell.

**Figure 6.**
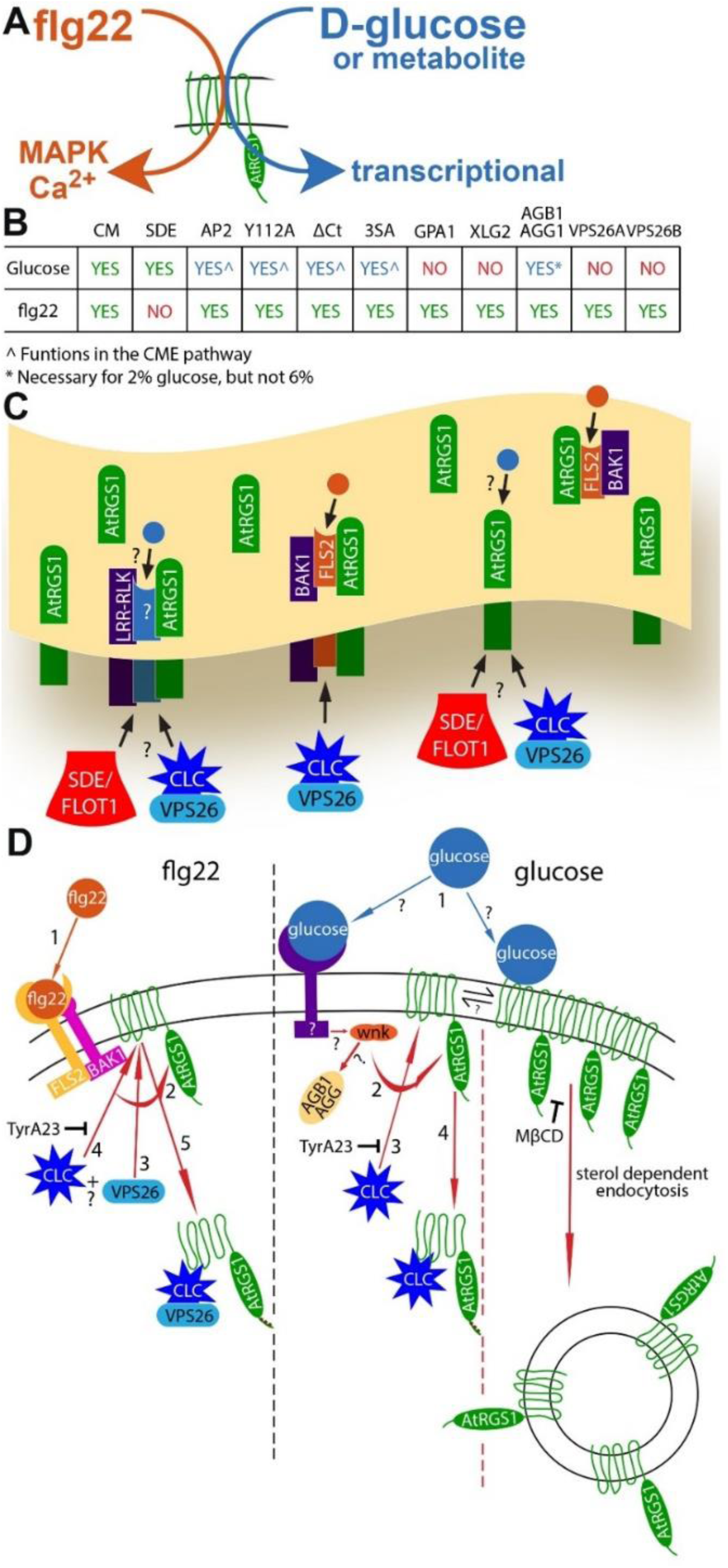
Component overview of flg22- and D-glucose-biased signaling. **(A)** Simple model illustrating flg22 and glucose or metabolite input and the respective bias signaling output through AtRGS1, as well as a chart summary detailing the origins of endocytosis (CME and SDE), recognition motif and phosphorylation requirements, and individual proteins necessary for glucose- and flg22-induced endocytosis of AtRGS1. (**B**) Membrane overview illustrating proposed AtRGS1 microdomain clusters with common RLK neighbors. flg22 (orange circle) binds to FLS2 to initiate signaling through AtRGS1. The mechanism of glucose (blue circle) perception is unknown as indicated by the question marks. After ligand perception, SDE or CME of AtRGS1 occurs to permit downstream signaling. (**C**) A diagram of the individual components involved in the mechanism of endocytosis initiated by flg22 and glucose or metabolite. Numbers indicate the order of operations. For flg22: (1) perception of ligand, (2) phosphorylation of AtRGS1 by a RLK, (3 and 4) binding of clathrin complex and/or VPS26 in an unknown order, and (5) internalization of AtRGS1. For glucose or metabolite: (1) ligand perception by a RLK or direct interaction with AtRGS1, (2) receptor interaction with WNK kinase (in the case of SDE, #2 indicates immediate endocytosis because other key components of the pathway are unknown), (3) phosphorylation of AtRGS1 by WNK kinases, (4) binding of clathrin complex to AtRGS1, and (5) internalization of AtRGS1. TyrA23 is shown inhibiting CME in both pathways, while MβCD is shown inhibiting AtRGS1 microdomain formation at the membrane.

The two mechanisms of AtRGS1 endocytosis induced by D-glucose or its metabolites implies that this signal-dependent pool of AtRGS1 is bipartite, prompting the question of an equilibrium or steady state between the populations. The evidence so far suggests that it is not. If these AtRGS1 pools were dynamically exchanging within the membrane, we would expect to induce internalization of nearly all membrane bound AtRGS1 by inhibiting one mechanism of endocytosis thereby causing a shift entirely to the other. For example, inhibiting CME with TyrA23 would force AtRGS1 endocytosis entirely to the SDE mode, however this was not observed. Inhibiting one mechanism of endocytosis only partly reduced AtRGS1 internalization by half, indicating the AtRGS1 populations may be physically isolated and static in the cell membrane (Figure 6C). Some AtRGS1 proteins may be grouped into so-called microdomains or clusters surrounded by receptor/co-receptor RLK proteins while other AtRGS1 proteins may be distributed throughout the membrane without common-neighbor RLKs (Figure 6C).

Similarly, the question exists whether an equilibrium between the flg22-mediated and D-glucose-mediated CME pools exists and the conclusion is again that an equilibrium is not likely, consistent with a genuine bias in signaling. To clarify what we define as the “CME pools” here, they are the portion of either the flg22- or D-glucose-activated AtRGS1 population that is inhibited by TyrA23, by loss of the tyrosine binding motif, and by loss of the AP2µ subunit of the clathrin complex (Figure 3). While the two pools share these properties, they do not share the requirement of the candidate adapter, VPS26 monomer or dimer (Figure 5). Moreover, when the AtRGS1 pool at the plasma membrane is depleted with D-glucose, there is no effect on the amplitude of flg22 activation (Figure 2).

The two mechanisms in the composite glucose-induced AtRGS1 endocytosis may be a result of two distinct mechanisms of sugar perception: one through direct interaction of AtRGS1 with a sugar or sugar metabolite and the other through sugar binding to a RLK in the membrane similar to flg22::FLS2 binding (Figure 6D).

To compare and contrast the two ligand-biased trajectories, the ordered steps of signal transduction from ligand perception (**step 1**) to internalization of AtRGS1 (**step 5**) are enumerated in Figure 7D to illustrate mechanistically what the present work revealed about each origin of endocytosis including phosphorylation (flg22- **step 2**/glucose- **step 2**), binding of the candidate adaptor VPS26 (flg22- **step 3**), formation of clathrin coated vesicles (flg22- **step 4**/ D-glucose- **step 3**), and finally endocytosis of AtRGS1 either via clathrin coated vesicle (flg22- **step 5/** D-glucose **step 4**) or sterol dependent rafts (glucose). If D-glucose or a metabolite is perceived by two distinct sensing mechanisms, each mechanism may operate exclusively through one origin of endocytosis with a unique core of G protein and internalization signaling components. Interestingly, within the glucose model, a system bias may exist favoring one origin of endocytosis that results from architectural differences in the membrane surrounding AtRGS1. A high density of D-glucose-binding RLKs may favor D-glucose-induced CME of AtRGS1 over the SDE origin.

The division of G protein involvement in D-glucose-induced CME and SDE challenges previously published reports on the necessity for subunits of the G protein heterotrimer in the full range of D-glucose signaling through AtRGS1; specifically shown was the complete abrogation of AtRGS1 endocytosis in the Gβ null mutant and partial reduction of D-glucose-induced endocytosis with the loss of Gα ^29^. Our results confirm that Gβ is required for AtRGS1 endocytosis, but only in a low dose scenario, 2% (∼110 nM). We show no requirement for Gβ or Gα at the higher 6% (∼330nm) glucose concentration. It may be that higher sugar concentrations, those typically found at or around vascular unloading areas, illicit a quicker signaling response than low dose sugar likely found in or near epidermal cells where we quantified AtRGS1 internalization. The significance of sugar signaling with regard to cell growth, division, and maintenance may necessitate multiple dose-dependent mechanisms of signal transduction encoded in different origins of endocytosis and the individual G protein associated components.

Both flg22- and D-glucose outputs require AtRGS1 ^30,41,58,90^ and an intact heterotrimeric G protein complex (Figure 4) but as discussed for D-glucose responsiveness, the outputs depends on both the concentration and duration of D-glucose. We previously designated this non-threshold-based activation phenomenon as Dose-Duration Reciprocity (**DDR**) where a low dose of D-glucose for a long period reaches the same output amplitude as a high dose presented as a pulse ^2^. The proposed mechanism is recruitment of WNK8 and WNK1 to the membrane by the Gβγ dimer AGB1/AGG as a function of DDR to phosphorylate AtRGS1 for endocytosis. However, our higher resolution analyses here challenge some aspects of that original model and provide deeper mechanistic insight. AGB1 is essential for low D-glucose DDR but is not required for high-glucose DDR. WNK8 kinase is required as previously shown but WNK1 is shown to not be required at longer, low-glucose treatments (6 hours) in contrast to previously published data ^2^. This discrepancy is most likely due to differences in expression level between the two studies (transient for ref. 2 vs. stable here) because it is known that glucose responsiveness is sensitive to the pool size of AtRGS1 ^111^.

Besides the 2 visual and 2 non-visual (aka β-) arrestins, the arrestin super family in humans also has 6 α-arrestin members. These α–arrestins have diverse functions including trafficking of non-GPCRs ^112^ and may be closest to the ancestral arrestin from which the visual/non-visual arrestins evolved because they are found in stromenopiles basal to the opistokont lineage containing metazoans and fungi ^92^. Therefore, we asked whether Plantae contains α–arrestin homologs and are thus the true adaptors for trafficking plant 7TM proteins. While we found clear α–arrestin orthologs in the 4-cell alga *Tetrabaena* (PNH03120) and in the green alga *Chara* (<u>GBG71752</u>), both basal to land plants, using *Mus* ARRDC3/TLIMP (<u>BAC65781</u>) as the search query, no hits were found in Arabidopsis (eudicot), rice (monocot), *Amborella* (base of angiosperms), and *Physcomitrella* (basal to higher land plants). This suggests that α–arrestins were lost in Plantae and this loss possibly allowed expansion of the related VPS26 family to evolve another function ^92^.

Given the fact that many GPCR-containing organisms divergent to humans lack arrestin proteins, another important finding from this study is the suggestion that the retromer subunits AtVPS26A and B moonlight as arrestin-like adaptors for phosphorylation-induced, clathrin-dependent AtRGS1 endocytosis in Arabidopsis. Could this function be ubiquitous? Even in humans, some GPCR endocytosis occurs independently of β-arrestins yet still requires C-terminal phosphorylation, a tyrosine motif, AP2, and clathrin (e.g. ^113,114^) raising the possibility that VPS26 orthologs in humans may serve the exact adaptor function for a subset of GPCRs.

This study focuses on biased signaling launched from different architectures of two AtRGS1-centered, physically-distinct pathways. We show that phosphorylation by different sets of kinases encodes this bias. However, we have not address how the architecture is established or maintained but we speculate that this too is based on a phosphorylation bar code on the AtRGS1/G protein complex. Therefore, establishing the dynamics of both the pre- and post-signaling phosphorylation bar codes is important for our understanding of biased system signaling.

Why both cytoplasmic and extracellular glucose pools are monitored by the plant cell remains unclear. We speculate that the extracellular pool of glucose is far more dynamic than the cytoplasmic pool due to its rapid metabolic flux in the cytoplasm compared to the apoplast. Therefore, an extracellular glucose detection system may be more appropriate for monitoring sugars produced by photosynthesis for real time information. This is consistent with the recent finding that AtRGS1 is important for detecting fluctuations in CO_2_-fixed sugar in the minute time range over the diel cycle ^59,90^.

It is conceivable that this biased system sits at the crux of the “defense vs. growth trade-off” dilemma that plants face. Specifically, pathogen attack compels the plant to shift its utilization of fixed sugars from building cell walls to synthesizing defense molecules ^115^. The AtRGS1/G protein complex may be the fulcrum for this balance because AtRGS1 is involved in detecting fixed sugars ^59,116^, establishing cell wall composition ^117–119^ and serving as a sentinel in innate immunity ^33^.

In conclusion, our data provide evidence for biased-system signaling through AtRGS1 and introduces a previously unknown arrestin-like adaptor. We introduce the importance of system architecture as it relates to bias in G protein complex signaling. We show that while phosphorylation is a requisite for AtRGS1 internalization, the direct phosphorylation of di-serines on AtRGS1 is not the signal discriminator. This likely comes from RLK interaction with other components in the signaling pathway. Subsequent analysis of protein clustering in the membrane and the phosphorylation barcode are necessary to understand the full extent of the system bias as it relates to signal transduction through AtRGS1.

## Methods

### Chemicals

Methyl-β-cyclodextrin was purchased from Frontier Scientific, tyrphostin A51 and turanose were purchased from Sigma-Aldrich, and tyrphostin A23 was purchased from Santa Cruz Biotechnology. All chemicals were indicated by the vendors to be >98% pure.

### Plant growth conditions

Wildtype and mutant Arabidopsis seeds (all Col-0 ecotype) were surface sterilized with 80% EtOH for 10 seconds while vortexing followed by a 10-second vortex with 30% bleach. Seeds were subsequently washed 3X with ddH_2_0 and suspended in 12 well cell culture plates with ¼ MS with no sugar at pH 5.7 with 10-12 seeds per well. Plates were wrapped in aluminum foil and cold-treated at 4°C for 2 days followed by a 2-hour light treatment to induce germination. After light treatment, plates were again wrapped in aluminum foil and placed on a horizontal shaker at ambient temperature for 5 days before imaging.

### AtRGS1 internalization assay, proxy for G protein activation

AtRGS1-YFP internalization was induced with D-glucose and flg22 as described ^2,29,32^. Briefly, Arabidopsis seeds expressing 35S:AtRGS1-YFP were sterilized and then sown on 1-mL liquid 1/4 X Murashige and Skoog (MS) medium without sugar in 12-well plates and stratified at 5°C for 2 days, followed by 2 hours light, then grown in darkness at RT for 3-5 days. For optimal results, the plates were kept in darkness but moved to the microscope room on the third day to acclimate. 6% D-glucose or 1µm flg22 were applied to seedlings for 30 and 10 minutes respectively before imaging. Image acquisition was done on either a Zeiss LSM710 (Zeiss Microscopy, Oberkochen, Germany) with a C-Apochromat 40X 1.2NA water immersion objective (for figures 2A-2I and figure 3A-3D) or a Zeiss LSM880 with a C-Apochromat 40x/1.2NA water immersion objective (for figure 2K-2P, 3E-3K, and S3). YFP excitation was at 514nm and emission collection 525-565nm. Emission collection on the LSM880 was done with a GaAsP detector. For RGS internalization assays a z-stack series was acquired at 0.5µm intervals between images. Image processing and RGS internalization measurements were done with the Fiji distribution of ImageJ ^120^ as described by Urano *et al* ^29^ with the following modification: Internalized YFP fluorescence was measured and subtracted from total YFP fluorescence of individual cells as opposed to total fluorescence of the hypocotyl image as stated in Urano *et al*. Images were acquired on the hypocotyl epidermis 2-4 mm below the cotyledons of seedlings treated with water, glucose, and flg22 in addition to the pharmacological inhibitors. Seedling exposure to light was minimized as much as is practical while imaging to avoid light induced internalization of AtRGS1. Statistical analysis was performed using analysis of variance with n=number of cells measured. Differences in basal and treatment induced levels of RGS1-YFP internalization are a result of switching image acquisition from the LSM710 (Figures 1, 2, & 4) to the LSM880 (Figure 3) to improve image resolution and accuracy of internalization measurements.

### Pharmacological inhibition of RGS internalization

AtRGS1 internalization was inhibited with TyrA23 and MβCD under the following conditions. TyrA23 was applied to 3-day old seedlings for a pre-incubation period of 60 minutes at specified concentrations. Following the pre-incubation period, a combination of TyrA23 and 6% D-Glucose were applied to the seedlings for 30 minutes immediately followed by image acquisition. In the case of flg22, TyrA23 and 1µm flg22 were applied to the seedling for 10 minutes following the pre-incubation period. For MβCD, the pre-incubation period was 45 minutes at specified concentrations. When both inhibitors were simultaneously applied, pre-incubation was 60 minutes

### TIRF imaging and area/speed measurements of AtRGS1-GFP

Arabidopsis expressing 35S-RGS1-GFP were grown as mentioned in the plant growth section. 5-day-old seedlings were transferred to a solution of either 6% D-glucose or 1µm flg22 and imaged at 5, 10, and 15 minutes while immersed in the ligand solution. Imaging was performed on a Nikon Ti Eclipse with SR Apo TIRF 100x lens (NA 1.5, WD 120µm). GFP excitation occurred at 488nm and emission collection at 515-555nm with an Andor iXon3 EMCCD camera. 60 second time-lapse imaging was initiated at the beginning of each time point with 200ms acquisition speed. Time-lapse sequences were normalized for fluorescence over time using IMARIS (v9.2.2, Bitplane AG, Zurich, Switzerland). The IMARIS Surface feature was used to track and calculate the area and speed of individually identifiable AtRGS1 proteins/clusters (labeled as tracks in IMARIS) over time. The average speed and area of each unique track for a 30 second interval between 5:15-5:45 or 15:15-15:45 was calculated using a script in Matlab (Supplemental code).

### Synthesis of [^14^C] isomaltulose

Sucrose isomerase (SI) from *Pantoea dispersa* UQ68J (GenBank AY223549) was cloned into expression vector pET24b (Novagen) first by PCR of genomic DNA using the following PCR forward primer 5’-GGA TCC AAC AAT GGC AAC GAA TAT ACA AAA GTC C-3’ which included a *BamHI* restriction site and a start codon; reverse primer 5’-ATA GGT ACC TCA GTT CAG CTT ATA GAT CCC-3’ which included a *KpnI* restriction site and a stop codon.

Expression was performed using *E. coli* BL21(DE3) (Novagen), 37°C, 225 rpm. When the optical density at 600nm reached 1.00, isopropyl-D-thiogalactopyranoside was added to a final concentration of 0.5 mM for induction. The incubation of the culture was continued for another 3 h at 28°C. Cells were harvested by centrifugation (3,000 ×g, 4°C, 10 min), resuspended in 50 mM Tris-HCl (pH 8.0)-2 mM EDTA, and then re-centrifuged. The cell pellet was immediately frozen in liquid nitrogen and stored at −75°C. Cells were suspended in extraction buffer (20 mM Tris-HCl (pH 7.4), 200 mM NaCl, 1 mM EDTA, 1 mM azide, 10 mM β-mercaptoethanol) and then lysed by sonication (nine 15-s pulses at 50 W with a Branson Sonifier 450 microprobe), centrifuged (10,000 ×g, 4°C, 10 min), and filtered through a 0.45-µm-pore-size membrane (Gelman Acrodisc). The pET24b vector introduced a carboxy-terminal six-His tag into expressed proteins, which were purified by adsorption to nitrilotriacetic acid (NTA) agarose (QIAGEN) and elution with 25 mM NaH2PO4-150 mM NaCl-125 mM imidazole buffer (pH 8.0) by following the manufacturer’s instructions. The purity of SI proteins was verified by SDS-PAGE as a single band on Coomassie Blue R-250 staining.

[^14^C] isomaltulose was prepared using 1.48MBq [U-^14^C]Sucrose (Amersham, UK) in 200 µl water with 3% ethanol (equals to 0.3379 mM) was reacted with 30 µl purified UQ68J SI for at 30 °C 60 min. The converted [^14^C] isomatulose concentration by UQ68J SI was estimated by three-replicate parallel conversions of unlabelled sucrose (S7903, Sigma) in the same concentration of 0.3379 mM with 3% Ethanol by the same enzyme. BioLC DX600 (Dionex, USA) determinations showed 84.0±0.106% (Mean±SE) was converted into isomaltulose, 3.5±0.197% into trehalulose; into the by-products of glucose and fructose were 4.6±0.072% and 7.8±0.237% respectively and there was no sucrose left after the reaction was stopped (Figure 1A).

### Sugar uptake assay

One-week-old seedlings, grown on a filter disc overlaying 1/2X Murashige and Skoog Basal Salts, 0.7% phytogel 23°C, pH 5.8, 8h/d of 100 mole/m2/s1, were lifted off the plate and overlaid 6 mL of water containing approx. 25,000 cpm of [^14^C] sugars as indicated. The specific activity of the sugars was 12GBq/mmol. At the indicated times, triplicate sets of 10 seedling were gently rinsed and placed in a 1.7-mL microfuge tube with 1 mL of scintillation fluid (Perkin Elmer Inc) and radioactivity was quantitated by liquid scintillation counting. CPM from time zero (typically 80-150 cpm) was subtracted from the average of the 3 samples. The CPM for [^14^C] isomaltulose uptake into seedlings at each time was corrected for its 84% purity. The experiment as shown was repeated once with the same result. [^14^C] glucose uptake was repeated 4 times.

### Live cell imaging of MAPK reporter (SOMA) lines

Detached etiolated hypocotyls were prepared for imaging on the confocal microscope using the HybriWell™ method as previously described ^71,121^. A hypocotyl from a dark grown 5-day-old seedling was placed on top of the droplet, and a HybriWell™ (Grace BioLabs, http://gracebio.com/, cat. no. 611102) was gently placed on the cover slide with the hypocotyl in the center to form a 150-µm deep imaging chamber with a volume of 30 μl. Ultrapure water (300 μl) was injected through one of the HybriWell™ ports using a pipettor to fill the 30-μl chamber with water and to expel any air bubbles. A 200-μl droplet of ultrapure water was then placed on one of the ports to prevent the chamber from drying out. The HybriWells containing the mounted hypocotyls were then placed in covered Petri dishes and equilibrated by incubating at 20–23°C under constant light for 6–8 hours prior to imaging.

Confocal microscopy was performed using a Zeiss LSM 710 with a C-Apochromat 40x/1.20 water immersion objective lens. Samples were excited at 458nm with 3% power, and emission was measured between 463 and 517nm for Turquoise GL and between 534 and 570nm for YPet. Z-stacks were collected every 2 min with an optical slice thickness of 1.2 μm. Chemical treatments were added to the samples during imaging by pipetting 200 μl of solution containing the treatment onto one port of the HybriWell. For experiments involving tyrphostin A23 and tyrphostin A51, hypocotyls were pretreated with 50 μM of these compounds for 30 min prior to imaging.

Post-processing of the raw image data was performed using Fiji ^120^. The ‘Z-projection’ function was performed on an image stack using the ‘Max Intensity’ setting. The resulting projection was then separated into two images, one for the Turquoise GL emission channel and one for the YPet emission channel. The ‘Subtract Background’ function was performed on both images, with the ‘rolling-ball radius’ set as the default 50 pixels. A mask was then created from the YPet channel using the ‘Convert to Mask’ function. The background subtracted YPet and Turquoise GL images were then converted into 32-bit images. These 32-bit images were then multiplied by the Mask file. The resulting YPet image was divided by the resulting Turquoise GL image using the ‘Image Calculator’ function to create a ratio image representing the ratio of YPet to Turquoise emission. Finally, the ‘Threshold’ function was performed using the default values, with the ‘NaN background’ option enabled. The ‘Fire’ lookup table was then applied to the final ratio image. To measure the ratio of YPet to Turquoise GL emission, a region of interest (ROI) was selected within the ratio image using Fiji and the average ratio value within that ROI was then measured.

### Live cell Ca^++^ imaging with R-GECO1

5-day-old, etiolated hypocotyls expressing R-GECO1 calcium reporter were grown in aqueous media containing ¼ MS. Hypocotyls were excised and mounted in HybriWells 6-8 hours prior to imaging with a Zeiss LSM710 confocal laser scanning microscope equipped with a C-Apochromat 40×/1.20 water immersion objective. R-GECO1 was excited using 561nm laser with 7.0 % laser power, and emission was measured between 620 and 650 nm. Z-stacks were collected 2 min after chemical treatment with an optical slice thickness of 1.5μm. Chemical treatments were added to the samples during imaging by pipetting 200 μl of solution containing the treatment onto one port of the HybriWell. The digital images were analyzed with Fiji ^120^.

### Modeling AtVPS26

Five models designated atvps26a-1 through 5 were created by the MODELLER using the automodel script based on the human VPS26A template 2FAU. For evaluation and selection of the “best” model, we calculated the objective function (molpdf), Discrete Optimized Protein Energy (**DOPE**) score, GA341 assessment score and root mean square deviation (**RMSD**) between the model and the template. The best model has the lowest value of the molpdf and overall DOPE assessment scores. In addition, DOPE scores were calculated per-residue and the template and the five atvps26a models were compared using GNUPLOT (Figure S4C) and found to be dissimilar in only three positions (residue around 60, 240, and 260). The atvps26a2 model was selected based on the lowest, RMSD value (Figure S4D) and plotted DOPE per-residue score curve (Fig. S4E). With an RMSD of 0.17 Ȧ, atvps26a-2 differs from the template by less than the length of a C-C bond.

### Split firefly-luciferase assays

pCAMBIA/des/cLuc and pCAMBIA/des/nLuc ^122^ were used to generate the following plasmids: AtVPS26A-nLUC, AtVPS26B-nLUC, cLUC-AtVPS26A, cLUC-AtVPS26B, cLUC-AtVPS26like, cLUC-AtGPA1, AtAGB1-nLUC and AtRGS1-nLUC. pART27H-mCherry-AtAGG1 plasmid was obtained from Dr. Jose R Botella (University of Queensland, Brisbane, QLD, 4072, Australia). All plasmids were transformed into *A. tumefaciens* strain *GV3101*. nLUC and cLUC fusion partners were co-expressed in *N. benthamiana* leavesby agroinfiltration following protocols in ^123^. 48 hours after infiltration, 6mm Leaf discs were collected to 96-well plate and 40µl 0.4mM D-luciferin was added to each well. Luminescence was measured by a spectraMax L microplate reader (Molecular Devices).

### Yeast two-hybrid assays

Constructs used in yeast two-hybrid assays were pAS–RGS-J5 (amino acids 284–459), RGS-ΔCt (amino acids 284–416), RGS-ΔCtS405,406A, pACTGW–AtVPS26B and AtVPS26like. Y2H tested pairs were co-transformed into *AH109* yeast cells and plated on SD-LW media ^124^. Colonies were then inoculated into SD-LW liquid media and grew overnight at 30 °C. Then 1/10 diluted yeast cells were spotted on SD-LW, SD-LWH, and SD-LWH 10mM 3-AT (3-amino-1,2,4-triazole) plates. The plates were incubated at 30°C for 4 days and were imaged by gel documentation system (Axygen) under white light.

## Acknowledgements

We would like to thank Dr. N Phan for creating the AtRGS1^Y112A^ plasmid and transgenic lines and for experiments testing the involvement of adaptors in AtRGS1 endocytosis and data in Figure 1A and B. We thank Dr. V Gurevich, Dr. M Garcia-Marcos and Dr. CE Alvarez for helpful comments and Dr. K Ghusinga for assistance with writing MatLab code. We thank Dr. J Huang for help in the turanose experiment. This work was supported by grants from the NIGMS (GM065989) and NSF (MCB-1713880) to A.M.J. The Division of Chemical Sciences, Geosciences, and Biosciences, Office of Basic Energy Sciences of the US Department of Energy through the grant DE-FG02-05er15671 to A.M.J. funded the biochemical aspects of this project. The content is solely the responsibility of the authors and does not necessarily represent the official views of the National Institutes of Health.

## Author contributions

T.J.R-E., J.W., X.S., B.D., H.J., J.Y., M.T-O, F.L. and A.M.J. performed experiments and produced data used in the figures. T.J.R-E., J.W., X.S., and A.M.J. designed experiments, and analyzed results. P.K provided the MAPK reporters prior to its publication and guided those experiments. L.W. synthesized and purified [^14^C]isomaltulose. Y.T. performed sugar uptake experiments shown in Figure 1. T.J.R-E., J.W., and A.M.J wrote the paper.

## Supplemental Information

**Supplemental Figure S1.**
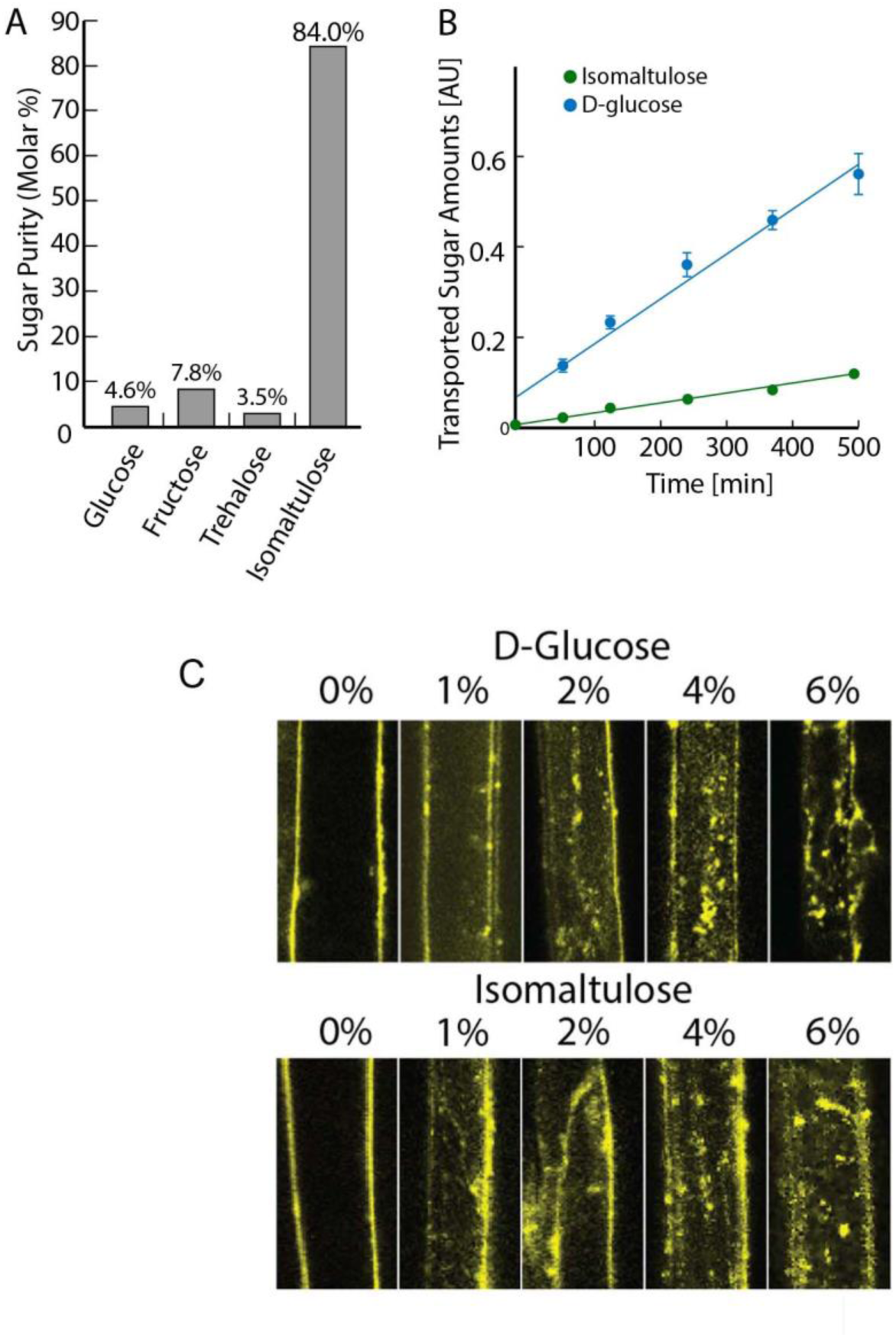
Isomaltulose is nearly impermeant in Arabidopsis. **A**. To test the permeability of Arabidopsis seedlings to isomaltulose, [^14^C] isomaltulose was synthesized and purified to 84% as described in the methods. Sugar analysis is as described and the results shown. **B**. Uptake of [^14^C] D-glucose and [^14^C] isomaltulose into Arabidopsis seedlings was tested. AU, Arbitrary Units as described. **C**. Isomaltulose and D-glucose induce rapid AtRGS1-YFP endocytosis in a dose-dependent manner. Images are representative from hypocotyl epidermal cells ectopically expressing AtRGS1 tagged with YFP.

**Supplemental Figure S2.**
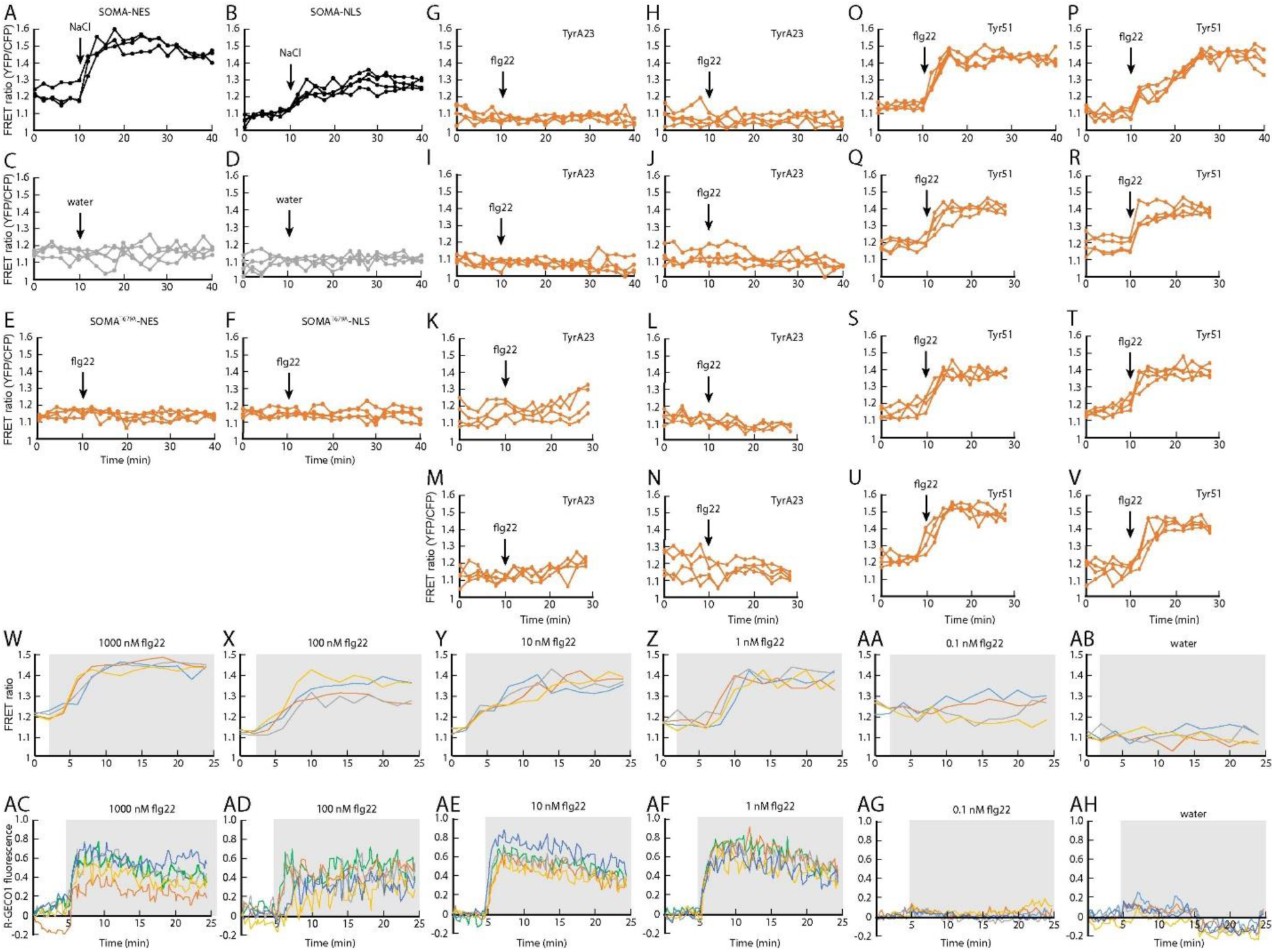
Positive and negative controls and dose response. Transgenic lines SOMA-NLS and SOMA-NES were analyzed before and after treatment with 150 mM NaCl as a positive control (**A**, **B**) and water (**C**, **D**). (**E**, **F**) Mutants of the transgenic lines SOMA^T679A^-NLS and SOMA^T679A^-NES were treated with 1 µM flg22 during the first 10 minutes of each experiment the samples. SOMA-NES lines were pretreated with 50 µM TyrA23 (**G**-**N**) or 50 µM TyrA51 (**O**-**V**) for 30 minutes prior to imaging. (**W-AB**) flg22 dose-response in SOMA-NES lines. (**AC**-**AH**) flg22 dose-response in R-GECO1 lines. Each graph represents 1 individual hypocotyl.

**Supplemental Figure S3.**
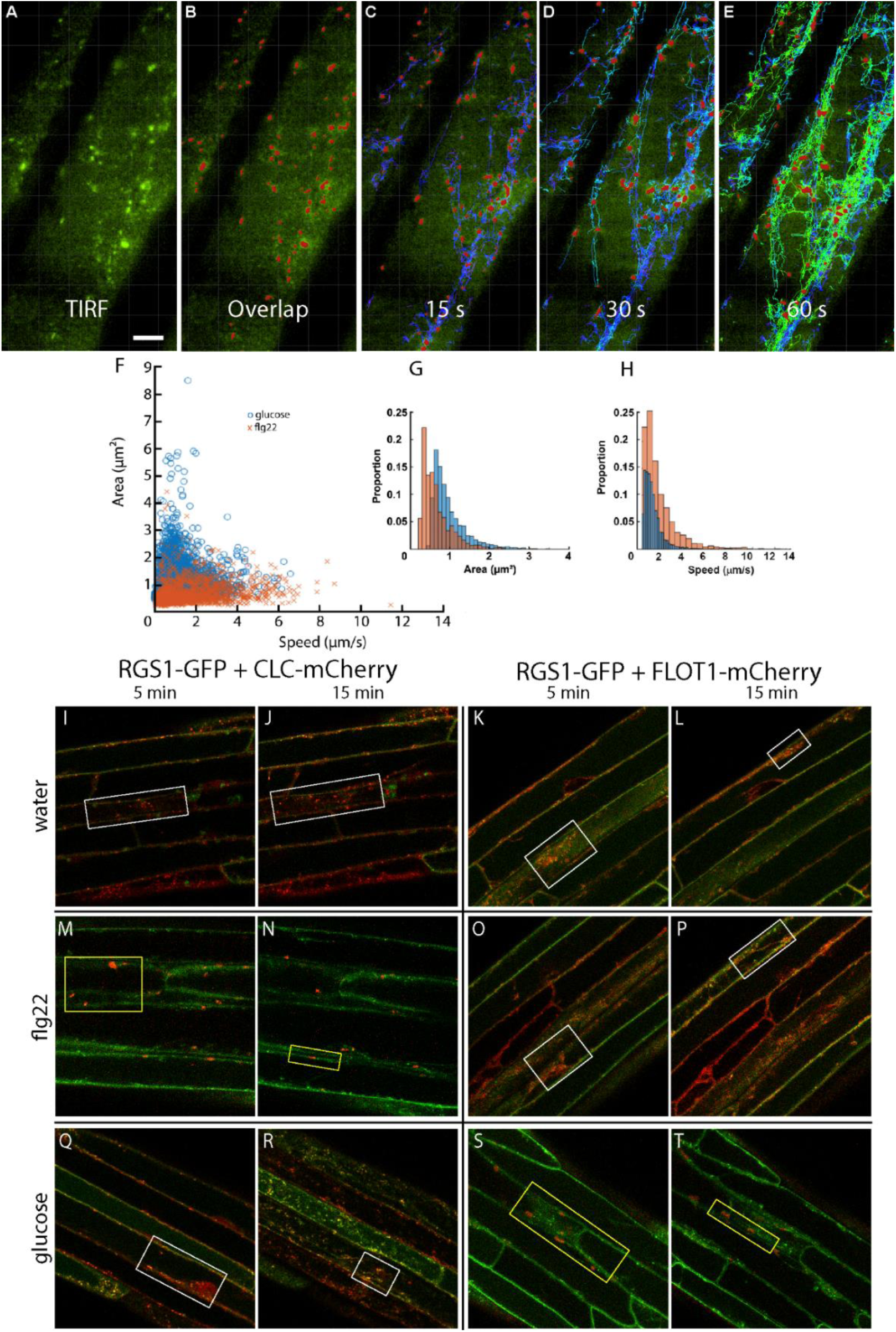
Tracking AtRGS1 after ligand addition. (**A-E**) IMARIS surface tracking showing (**A**) the original wide-field image acquired by TIRF, (**B**) the tracked spots imposed on the original image and tracked at (**C**) 15s, (**D**) 30s, and (**F**) 60s after initiating time-lapse imaging. (**F-H**) Tracking results showing the (**F**) average speed and area for AtRGS1-GFP particles after addition of (**x**) flg22 and (**o**) glucose at 15 minutes post ligand addition. (**G**) area and (**H**) speed of AtRGS1-GFP particles are broken out into proportion of total tracked particles at 15 minutes post ligand addition. (**I-T**) Original field of view confocal micrographs highlighting areas used to calculate Manders Overlap Coefficients from Figure 2K-P (white and yellow boxes) with the addition of 15 minute post ligand addition images.

**Figure S4.**
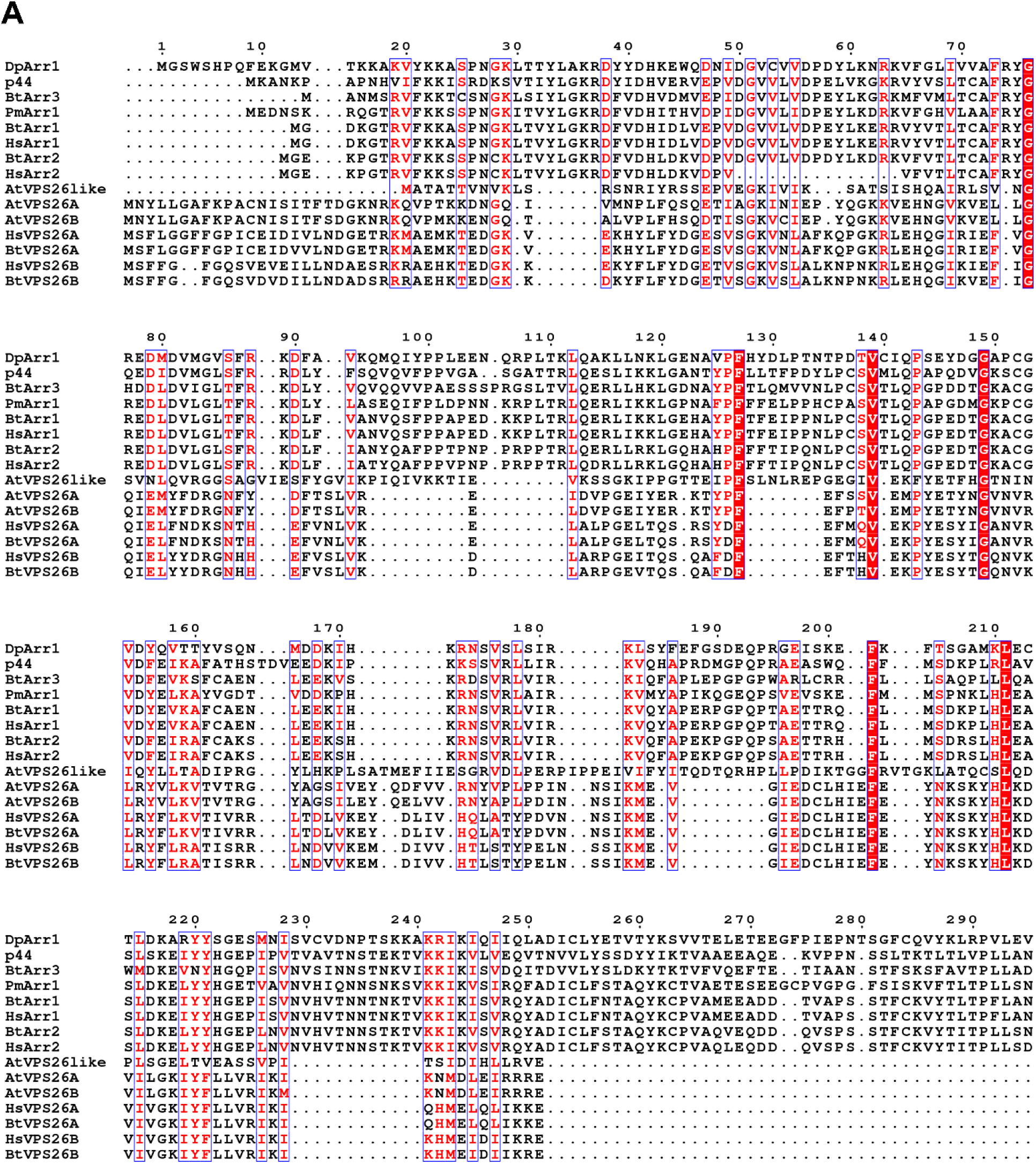
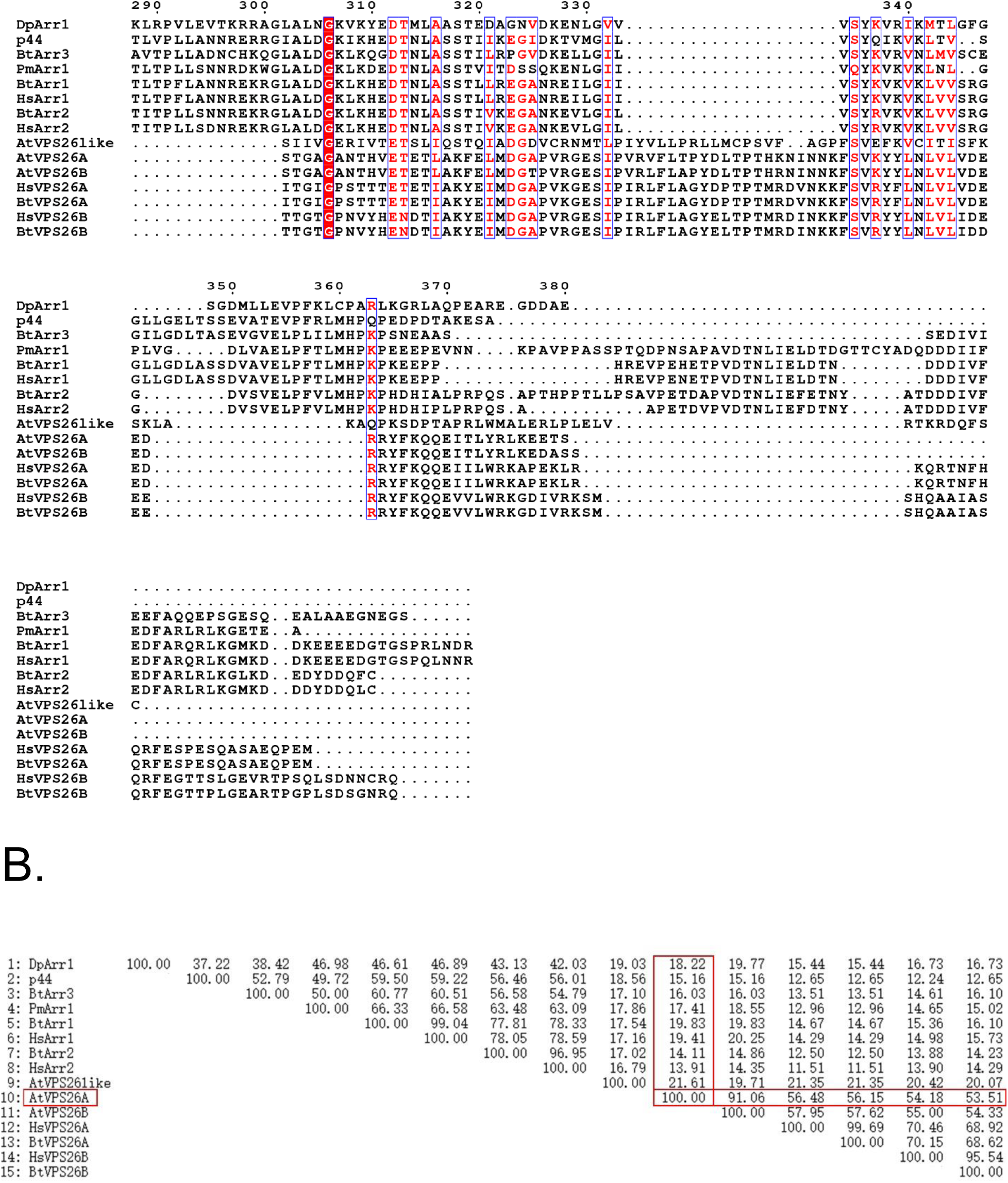
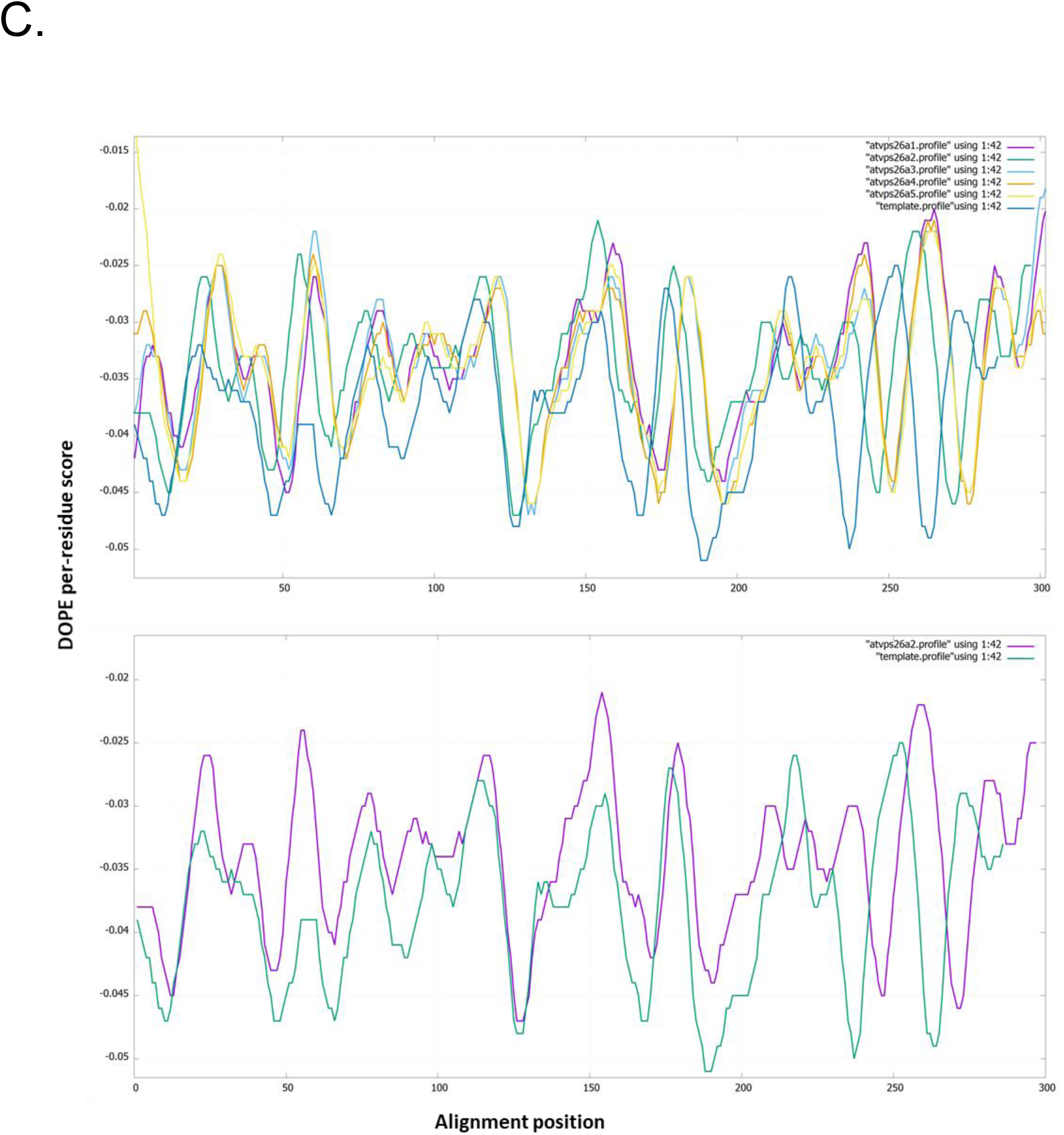
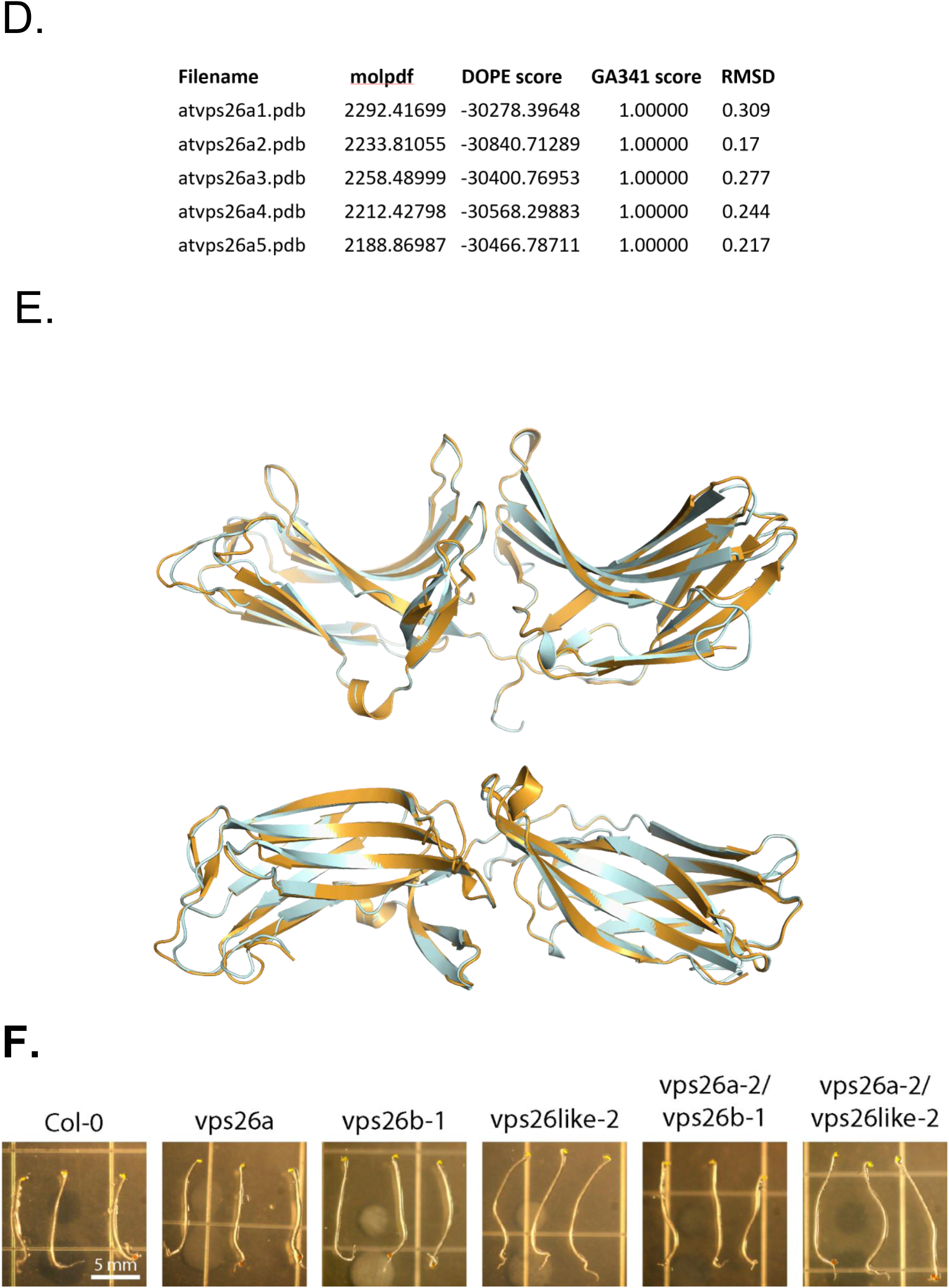
Supplemental data used to create the VPS26 model and comparison with arrestin structure described in main figure 4. (**A**) Homogeneous sequence alignment of VPS26 family (Arabidopsis VPS26a, VPS26b, VPS26like and Human VPS26A, VPS26B) with arrestin family (Human arrestin-1, arrestin-2 and bovine arrestin-1, arrestin-2, arrestin-3, squid arrestin-1, shrimp arrestin-1). (**B**) The identity matrix between arrestin family and VPS26 family. Clustal Omega https://www.ebi.ac.uk/Tools/msa/clustalo/ was used to generate the Multiple Sequence Alignment file and the identity matrix result. Then the ESPript 3.0 (Easy Sequencing in PostScript) which is a program which renders sequence similarities and secondary structure information from aligned sequences for analysis and publication purpose is used to generate the final alignment file http://espript.ibcp.fr/ESPript/ESPript/index.php. A percentage of equivalent residues is calculated per columns, considering physicochemical properties. A global score is calculated for all sequences by extracting all possible pairs of residues per columns, for the score greater than similarity GlobalScore (0.7), it was rendered as colored characters (red characters on a white background and white characters on a red background if residues are strictly conserved in the column) with blue frames. AtVPS26A has a high sequence identity of 91.06% with AtVPS26B and a 56.48% sequence identity with human VPS26A (shown in red box in figure B). AtVPS26A and the arrestin family share 14-20% sequence identity (shown in red box in the figure). Sequences: 1. DpArr1, squid arrestin-1; 2. p44, bovine arrestin-1 splice variant; 3. BtArr3, Bovine Arrestin-3; 4. PmArr1, shrimp arrestin-1; 5. BtArr1, Bovine Arrestin-1; 6. HsArr1, Human Arrestin-1; 7. BtArr2, Bovine Arrestin-2; 8. HsArr2, Human Arrestin-2; 9. AtVPS26like, Arabidopsis VPS26like; 10. AtVPS26A, Arabidopsis VPS26A; 11. AtVPS26B, Arabidopsis VPS26B; 12. HsVPS26A, human VPS26A; 13. BtVPS26A, bovine VPS26A; 14. HsVPS26B, human VPS26B; 15. BtVPS26B, bovine VPS26B.) (**C**) Model evaluation results of the 5 models of AtVPS26A. The MODELLER objective function (molpdf), DOPE assessment scores (Discrete Optimized Protein Energy, which is a statistical potential used to assess homology models in protein structure prediction), GA341 assessment score (range from 0.0 (worst) to 1.0 (native-like)) and RMSD (root-mean-square deviation, Å) with the template were calculated to evaluate the models. The “best” model is selected with the lowest value of the molpdf, DOPE score and RMSD value. The second model (atvps26a-2) was selected given the best DOPE score and RMSD value. (**D**) DOPE per residue score files of the 5 atvps26a models and the template. DOPE per residue score files of the five atvps26a models and the template human VPS26A[2FAU] were plotted using GNUPLOT which is a portable command-line driven graphing utility http://www.gnuplot.info/. Upper panel is the curves of the five atvps26a models and the template. The lower panel is the curve of the “best” model atvps26a-2 with the template. (**E**) 3D-structural alignment between model atvps26a-2 with human VPS26a template (PDB [2FAU]). atvps26a-2 model colored in pale-cyan and human VPS26A template colored in bright-orange using pymol. Upper panel: Side view. Lower panel: Top view. (**F**) Etiolated hypocotyls of Col-0 and VPS26 null mutants at 5 days old, the age used for experiments in Figure 5. Seedlings were germinated and grown in liquid MS for 5 days and transferred to solid agar plates for imaging. Scale bar 5mm. (**G**) Bifluorescence complementation (BiFC) of VPS26A and VPS26B showing a specific head-to-tail orientation requirement. Representative cells shown. n=5. Experiment repeated 2 times. (**H**) Yeast two-hybrid complementation between the cytoplasmic domain of AtRGS1 (RGS1-J5) and VPS26B. RGS1-J5 contains the linker between the 7TM domain and the RGS box, the RGS box and a C-terminal tail (CT). RGS1-ΔCT lacks the C-terminal tail which contains the phosphorylation cluster required for AtRGS1 endocytosis. RGS1-ΔCTS_405,406A_ lacks the CT and has two additional phosphosites mutated. –LW is the leucine/tryptophan dropout, -LWH is the leucine/tryptophan/histidine drop out media; 10 mM 3-AT indicates higher stringency by the addition of 10 mM 3-amino-1,2,4-triazole.

**Figure S5.**
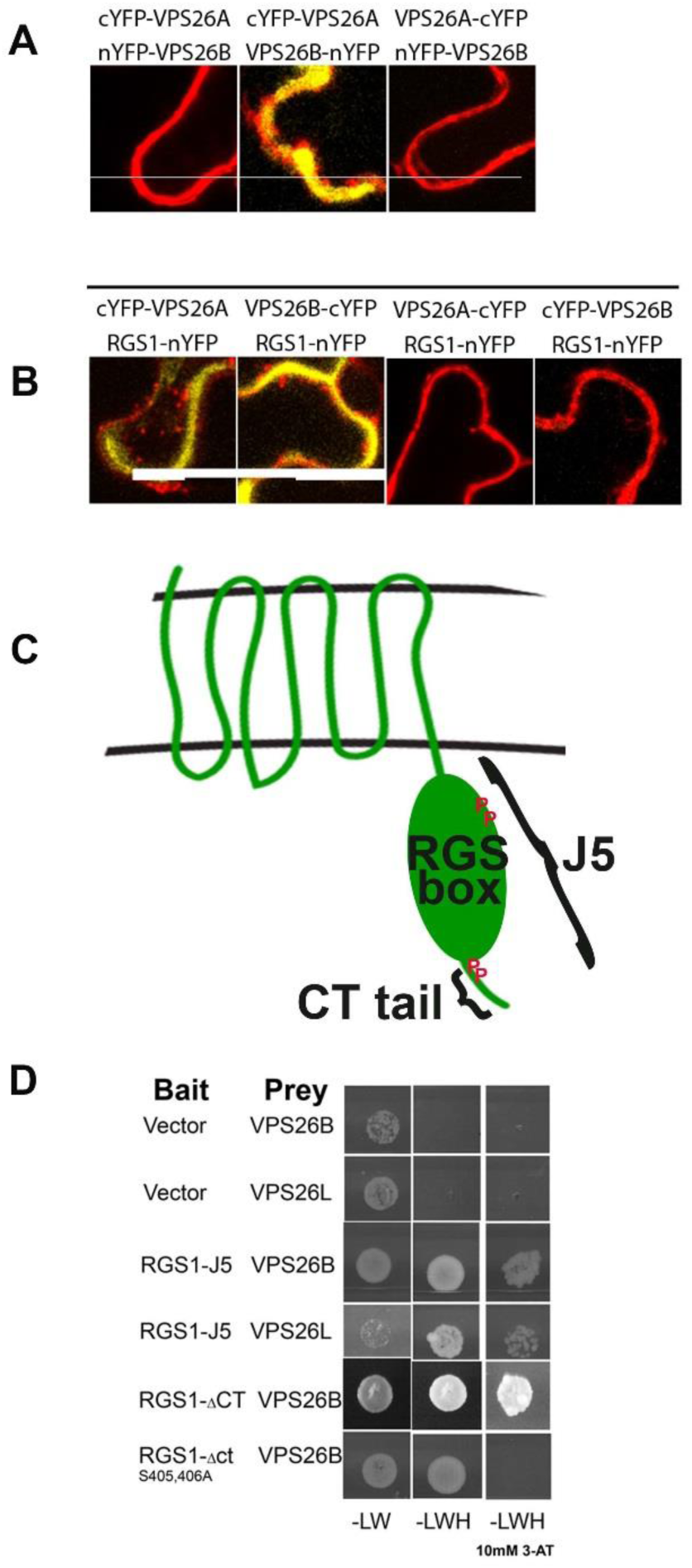
Supplemental dat for VPS26 dimerization, interaction with AtRGS1, and phosphoserine depence. **A)** Bifluorescence complementation (BiFC) of VPS26A and VPS26B showing a specific head-to-tail orientation requirement. Representative cells shown. n=5. Experiment repeated 2 times. **B**) Bifluorescence complementation (BiFC) of VPS26A and VPS26B interaction with AtRGS1 **C**) Definition of J5 cytoplasmic domain and the C-terminal (CT) tail. **D**) Yeast two-hybrid complementation between the cytoplasmic domain of AtRGS1 (RGS1-J5) and VPS26B. RGS1-J5 contains the linker between the 7TM domain and the RGS box, the RGS box and a C-terminal tail (CT). RGS1-ΔCT lacks the C-terminal tail which contains the phosphorylation cluster required for AtRGS1 endocytosis. RGS1-ΔCTS_405,406A_ lacks the CT and has two additional phosphosites mutated. –LW is the leucine/tryptophan dropout, -LWH is the leucine/tryptophan/histidine drop out media; 10 mM 3-AT indicates higher stringency by the addition of 10 mM 3-amino-1,2,4-triazole.

## Supplemental code

**Supplemental Code.**
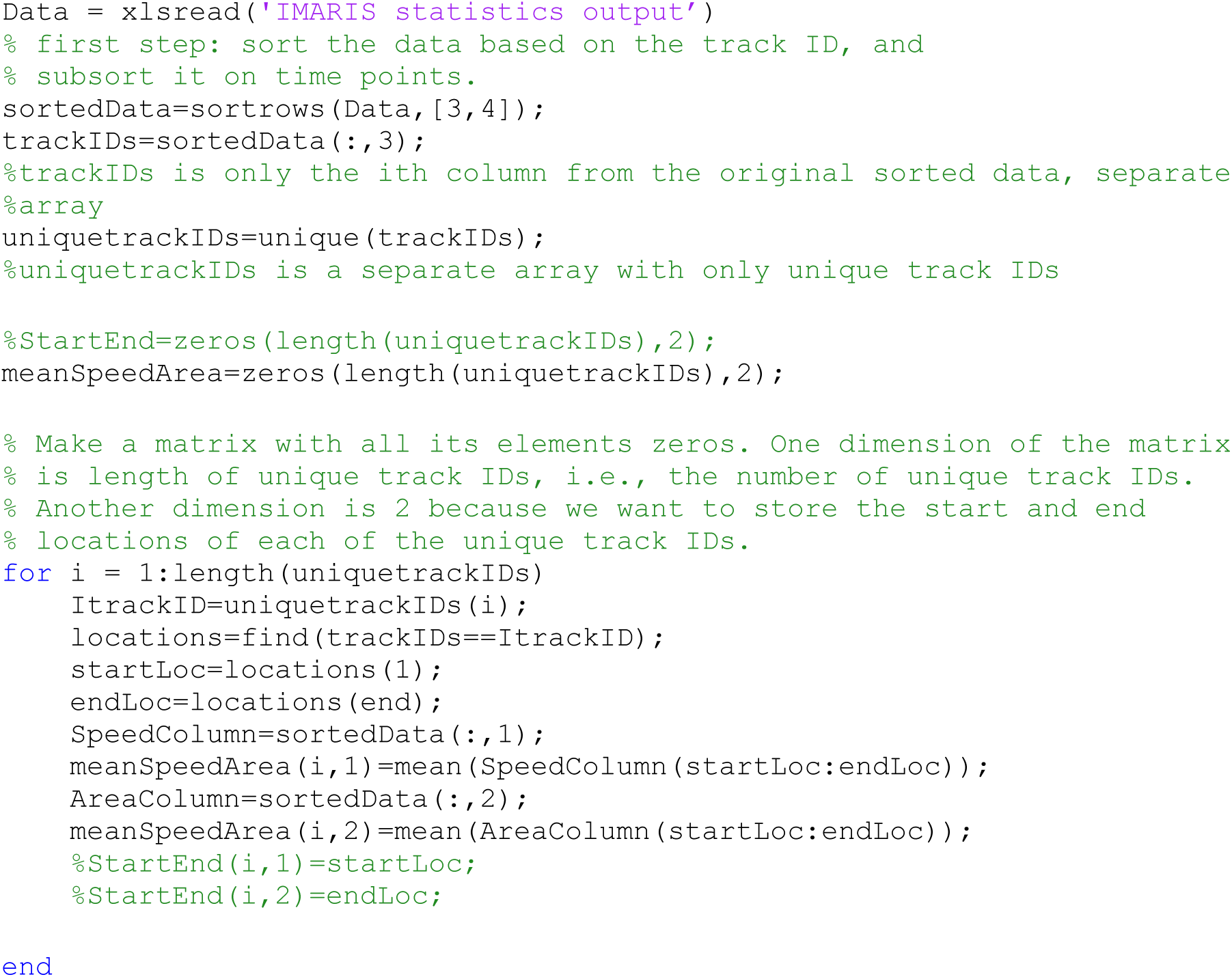
MatLab code for averaging unique tracked AtRGS1 particle speed and area. Speed and area data from IMARIS is exported in xls format and imported to MatLab for sorting. Each AtRGS1 tracked protein or cluster has a unique trackID that is sorted while maintaining association with speed and area at defined time points. We identified the start and end location for each unique trackID and created a matrix to store the data and subsequently find the mean for speed and area within our defined time points.

## Notes

#### Summary of Updates

Figures and summary have been updated.

